# Complex I Drives Glutamine-Dependent TCA Cycle to Support Viability of MYC^high^ Breast Cancer Cells

**DOI:** 10.1101/2025.05.27.656356

**Authors:** Johanna M. Anttila, Mariel Savelius, Juhi Somani, Daniel Nicorici, Pauliina M. Munne, Linda Id, Aino Peura, Antti O. Hiltunen, July Aung, Ryan Awadhpersad, Bina Prajapati, Martina Peltonen, Hanna Ala-Hongisto, Prson Gautam, Mika J. Välimäki, Topi A. Tervonen, Karina Šapovalovaitė, Raman Devarajan, María Victoria Ruiz Pérez, Minna Mutka, Panu Kovanen, Laura Niinikoski, Tuomo Meretoja, Johanna Mattson, Päivi Heikkilä, Krister Wennerberg, Marie Arsenian-Henriksson, Jukka Westermarck, Tero Aittokallio, Andrei Goga, Christopher B. Jackson, Anni I. Nieminen, Juha Klefström

**Affiliations:** Cancer Cell Circuitry Laboratory, Translational Cancer Medicine Research Program, Research Program Unit, Faculty of Medicine, University of Helsinki. PO Box 63 (Street address: Haartmaninkatu 8), 00014 University of Helsinki, Finland; Department of Biochemistry and Developmental Biology, Faculty of Medicine, University of Helsinki. Haartmaninkatu 8, 00290, Helsinki, Finland; Institute for Molecular Medicine Finland (FIMM), HiLIFE, University of Helsinki. Tukholmankatu 3, 00290, Helsinki, Finland; Department of Microbiology, Tumor and Cell Biology, Biomedicum, Karolinska Institutet, 17165, Solna, Sweden; Department of Pathology, HUSLAB and Haartman Institute, Helsinki University Central Hospital and University of Helsinki, Helsinki, Finland; Breast Surgery Unit, Comprehensive Cancer Center, Helsinki University Hospital and University of Helsinki, Helsinki, Finland; Department of Oncology, University of Helsinki & Helsinki University Hospital, Helsinki, Finland; Biotech Research and Innovation Centre (BRIC), University of Copenhagen, Copenhagen, Denmark; Turku Bioscience Centre, University of Turku and Åbo Akademi University, Tykistökatu 6A, Turku, 20520, Finland; Institute of Biomedicine, University of Turku, Turku, 20520, Finland; iCAN Digital Precision Cancer Medicine Flagship, University of Helsinki and Helsinki University Hospital, Finland; Institute for Cancer Research, Department of Cancer Genetics, Oslo University Hospital, Norway; Oslo Centre for Biostatistics and Epidemiology (OCBE), Faculty of Medicine, University of Oslo, Norway; Department of Cell & Tissue Biology and UCSF Helen Diller Comprehensive Cancer Center, University of California, San Francisco, 513 Parnassus Avenue, UCSF Campus Box 0512, San Francisco, CA 94143; Stem Cell and Metabolism Research Program, Faculty of Medicine, University of Helsinki; Finnish Cancer Institute, Helsinki, Finland; FICAN South, Helsinki University Hospital, Helsinki, Finland

## Abstract

In many cancers, stably elevated MYC levels drive persistent and concerted activation of cell growth promoting anabolic programs and the cell cycle in ways that are distinct from normal cells. Therefore, synthetic-lethal strategies to target MYC reprograming of these pathways may identify new selective anticancer therapies for the treatment of MYC^high^ tumors. Here, we identify enhanced mitochondrial respiration as a hallmark of MYC overexpressing cancer cells. Mitochondrial respiration sustains the TCA cycle by regenerating NAD^+^ through complex I-mediated oxidation of NADH. Metabolic carbon tracing analysis revealed that MYC shifts TCA cycle’s carbon source from glucose to glutamine. Inhibition of the glutamine-fueled TCA cycle using NAD^+^-depleting complex I inhibitors resulted in MYC-dependent synthetic lethality in breast cancer cells. In mouse models of MYC^high^ tumors, persistent inhibition of tumor growth was achieved through combined inhibition of complex I and glutaminolysis. Our results suggest that the high respiration rate observed in MYC^high^ cells supports glutamine carbon-enriched TCA cycle, rendering MYC^high^ tumors selectively vulnerable to inhibitors of mitochondrial respiration and glutaminolysis.

## INTRODUCTION

MYC is a basic helix-loop-helix leucine zipper (bHLH-Zip) transcription factor ^1^ and one of the most studied cancer driver genes. MYC protein is frequently expressed at elevated level in human cancer and the dominant presence of MYC^high^ cells in cancer associates with aggressive behavior, treatment resistance and unfavorable prognosis ^2,3^. For example, in breast cancer, about half of the tumors contain high proportions of MYC^high^ cells ^4,5^ and typically, these tumors represent the most aggressive subtypes such as triple-negative breast cancer (TNBC), Luminal B tumors, breast cancer metastases and drug-resistant tumors ^3,6–8^.

Elevated levels of MYC possess oncogenic activity, which deregulates a variety of cellular programs that control the cell cycle and proliferation, DNA replication and chromatin structure, metabolic pathways, cell adhesion, immunity, apoptosis, and many biosynthetic pathways ^9–13^. Some MYC reprogrammed pathways create specific vulnerabilities, which form rationale for therapeutic intervention via synthetic lethal strategies. Currently investigated targets, which upon inhibition predominantly or selectively harm MYC^high^ cells include, for example, cyclin-dependent kinases ^3^, anti-apoptotic BCL-2 family members ^5^, mammalian target of rapamycin complex 1/2 ^14^, aurora-B kinase ^15^ and polo-like kinase 1 ^16^. The principle of synthetic lethal strategies is to identify and target pathways and phenotypes highly enriched in the cancer cells – to direct the harmful effects of drugs on cancer cells while simultaneously sparing healthy cells to avoid drug-related adverse effects.

Some of the most selective MYC^high^ associated cellular phenotypes include metabolic dependencies, such as those rendering cancer cells reliant on specific nutrients. MYC overexpression has been shown to mediate dependence for example on non-essential amino acids like glutamine and cysteine ^17,18^. MYC-induced glutamine and cysteine dependence ^19^ is thought to arise from the onset of glutamine anaplerosis in MYC^high^ cells ^17^ as several reports have shown that MYC enhances glutaminolysis, which promotes influx of glutamine-derived α-ketoglutarate into the TCA cycle ^17,20,21^. In rapidly proliferating cells, many TCA cycle intermediates are being extracted for biosynthesis and the glutamine anaplerosis as a process replenishes the otherwise dwindling TCA cycle intermediates through provision of sufficient carbon supply for continued TCA cycle ^21,22^. The sustained TCA cycle consequently generates reducing equivalents (NADH, FADH2), which in turn drive the mitochondrial respiratory chain and generate biosynthetic precursors for lipids and nucleotides.

The MYC target gene expression signatures or “MYC activity” signatures may provide robust and quantitative metrics for evaluation of the MYC^high^ status of cancer cells to complement semi-quantitative and often antibody quality-dependent protein-based assays. Here, we identify a MYC signature that is highly correlated with MYC protein levels in a panel of 14 TNBC cell lines. Both transcriptomic and biochemical analyses revealed enhanced mitochondrial respiration in the MYC^high^ signature cells. The association between the MYC^high^ activity and enhanced respiration was corroborated in global analysis of the transcriptomes of cancer cell lines and tumor samples. Thus, high mitochondrial respiration appears as a universal MYC^high^ phenotype, present in TNBC and wide variety of cancer cell lines of different origins. The MYC^high^ TNBC cells were preferentially sensitive to metformin, a weak mitochondrial respiratory complex I inhibitor, which resulted in diminished proliferation together with apoptotic priming. Mechanistically, we show that MYC induces a metabolic reprogramming, shifting glutamine over glucose as the TCA cycle’s carbon source. Inhibition of complex I with metformin or ubiquinone (Q) reduction site blocking specific complex I inhibitors strongly repressed the glutaminolysis-dependent TCA cycle and selectively promoted death in MYC^high^ TNBC cells. In summary, we find that high MYC levels are coupled to high mitochondrial respiration rate and striking metabolic shift towards a glutamine-fueled TCA cycle in breast cancer. The MYC^high^ OXPHOS^high^ breast cancer phenotype marks a metabolic vulnerability targetable with low doses of complex I inhibitors alone or in combination with clinical stage inhibitors of glutamine metabolism.

## RESULTS

### Transcriptional signature that defines MYC^high^ TNBC cells

MYC activity signatures have been used, for example, to define subtypes of breast cancer with highest MYC activity for trials with MYC pathway directed drugs ^3,23^, to predict clinical outcomes among MYC associated cancers ^24,25^, and more recently, to define metabolic alterations that associate with MYC^high^ areas within mouse and human tumors ^26^. To define which published transcriptional MYC activity signatures correlate best with high levels of MYC protein expression, we determined the population mean MYC protein expression level by high content immunofluorescence (IF) analysis in 15 triple-negative breast cancer (TNBC) cell lines (Fig. 1A). TNBC represents 10-15% of all breast cancers and is an aggressive subtype with the lowest 5-year relative survival rate among different breast cancer subtypes ^6,27,28^. In addition to IF, bulk RNA-seq was performed in triplicates for the same cell lines to profile transcriptomic levels of MYC target genes. We compared the population mean MYC protein expression levels obtained via IF to MYC protein data from Western blot analysis ^5^ and mass spectrometry ^29^. We found significant Spearman correlation between the IF and Western blot results (Supplementary Fig. 1A) and between the IF and mass spectrometry assays (Supplementary Fig. 1B). Thus, our results demonstrate that cancer cell lines can be robustly and consistently categorized to MYC^high^ and MYC^low^ expressors via multiple protein-based assays.

**Fig. 1.**
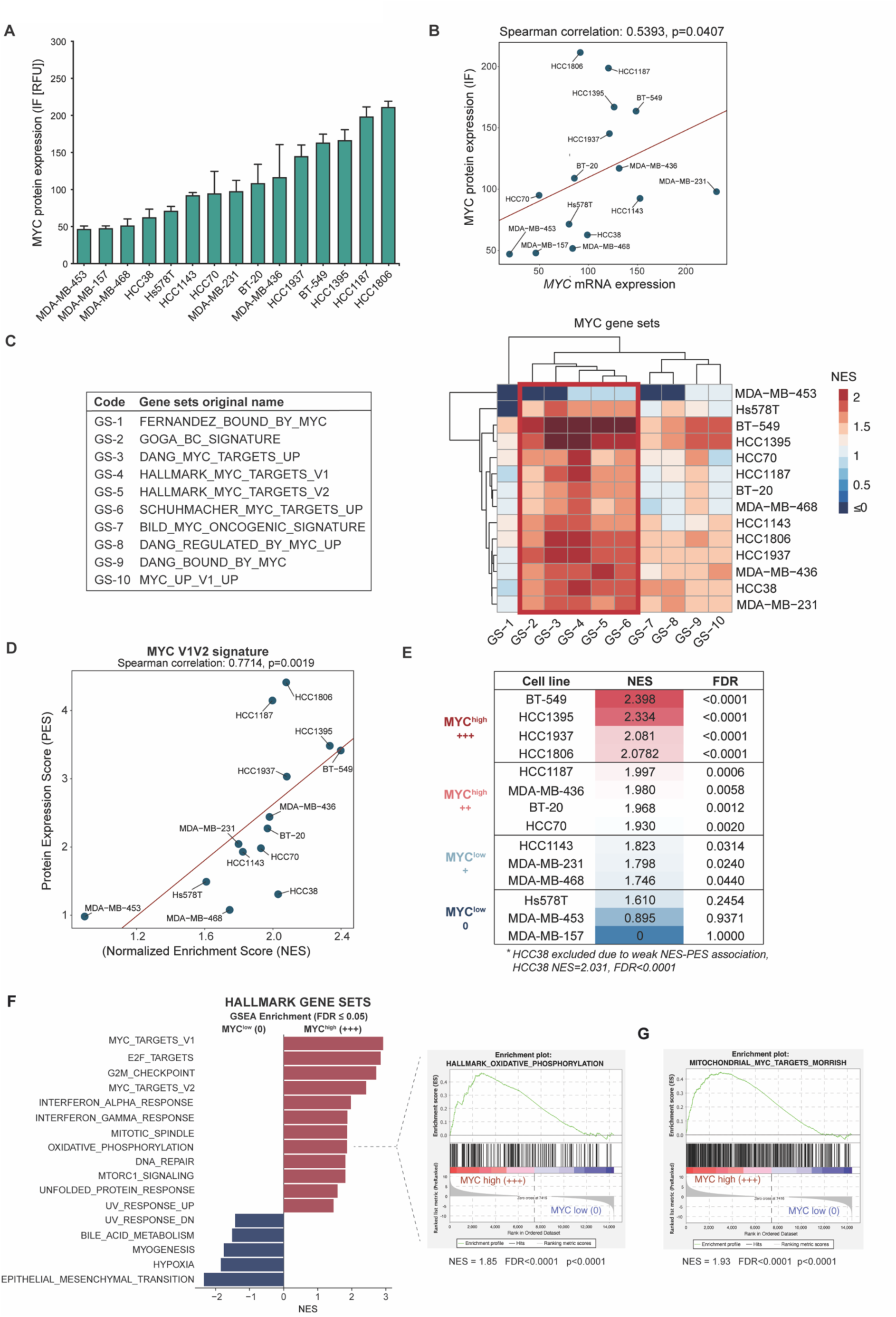
Stratification of TNBC cells to MYC^high^ and MYC^low^ groups. **A** MYC protein expression across 15 TNBC cell lines determined using high content immunofluorescence imaging. Data: mean +/− SD. **B** Spearman correlation analysis for the correlation of *MYC* mRNA expression (Transcripts Per Kilobase Million/TMP values) and MYC protein expression in TNBC cells. **C** Hierarchical clustering of MYC activity gene sets. Left panel: Nine MYC target gene sets were derived from the MSigDB collections and an additional MYC-related gene set referred as GOGA_BC_SIGNATURE (GS-2) (from ^13^) was included. Right panel: Hierarchical clustering of GSEA-based NES values for TNBC cell lines finds five similar gene sets (marked with red rectangle). MDA-MB-157 cell line was used as the reference cell line. **D** Spearman correlation between MYC V1V2 Normalized Enrichment Scores (NES) and Protein Expression Scores (PES) in TNBC cells. **E** Stratification of TNBC cells into MYC^high^ and MYC^low^ groups. TNBC cell lines were stratified into MYC^high^ and MYC^low^ groups according to the NES and FDR values from GSEA using MYC V1V2 gene set. The cut off for MYC^high^ +++ group was arbitrarily set as NES>2, FDR<0.0001 and for MYC^low^ 0 group NES<1.7, FDR>0.1. The NES and PES values correlated well except for HCC38, which was excluded from the analysis. **F** Differentially expressed Hallmark gene sets in MYC^high^ (+++) *versus* MYC^low^ (0) TNBC cell lines. **G** GSEA of mitochondrially localized MYC targets ^9^ in MYC^high^ (+++) *versus* MYC^low^ (0) TNBC cell lines.

Consistent with recent proteogenomic studies indicating overall low correlation between gene amplifications and corresponding protein expression ^30^, we did not find significant correlation between high *MYC* gene amplification status and a high level of MYC protein or mRNA (Supplementary Fig. 1C, D). However, *MYC* mRNA levels did correlate with the MYC^high^ protein status (Fig. 1B). To explore whether the published MYC target gene signatures correlate with the MYC^high^ protein status in TNBC cells, we used two gene set enrichment analysis tools: Gene Set Enrichment Analysis (GSEA) and Correlated Adjusted MEan RAnk gene set test (CAMERA). MDA-MB-157 cells expressed low MYC protein and mRNA levels as well as the lowest score for MYC target signatures (Molecular Signature Database MSigDB; Broad Institute). Therefore, MDA-MB-157 cells were used as a reference cell line in all subsequent analyses. From GSEA analysis, 10 MYC target gene sets (GS) were selected, and the normalized enrichment score (NES) was used to rank the 15 TNBC cell lines according to their inferred MYC activity status (Fig. 1C). We analysed the consistency of the GS-specific NES values across the 15 TNBC cell lines via hierarchical clustering, which revealed five gene sets with similar NES patterns (Fig. 1C, right panel).

We chose these five GSs for further analysis and correlated their NES values to the MYC Protein Expression Scores (PES) (Supplementary Fig. E-I). The best association between the NES and PES values was observed when GS-4 (HALLMARKS_MYC_TARGETS_V1) and GS-5 (HALLMARKS_MYC_TARGETS_V2) were combined into a larger MYC V1V2 set (Fig. 1D). Further, the CAMERA analysis confirmed the best PES correlation for MYC V1V2 (Supplementary Fig. 1J-L). The observed strong correlation between the MYC V1V2 gene set and the MYC PES status in two different analyses supported the use of MYC V1V2 set as a MYC activity-based biomarker for stratification of the TNBC cells into MYC^high^ and MYC^low^ groups (Fig. 1E). The highest MYC activity was attributed to a group of BT-549, HCC1395, HCC1937, HCC1806 and HCC38 cell lines (NES >2; FDR< 0.0001). However, HCC38 cell line was excluded from the MYC^high^ set as an outlier due to a low NES-PES correlation (Fig. 1D, Supplementary Fig. 1J). Lowest MYC scores (MYC^low^) were attributed to a group of Hs578T, MDA-MB-453 and MDA-MB-157 cells. Together, our analyses reveal the MYC V1V2 gene set as a strong RNA-based biomarker for MYC activity across multiple TNBC cell lines, which correlates with the MYC protein expression in TNBC cells and can be used to stratify breast cancer cell lines into MYC^high^ and MYC^low^ groups.

To explore molecular differences between the MYC^high^ and MYC^low^ TNBC groups, we first performed GSEA analysis on 50 hallmark molecular signatures in the Molecular Signatures Database (MSigDB; Broad Institute, ^31^. Notably, the hallmark oxidative phosphorylation set was among the significantly enriched gene sets in the MYC^high^ group (Fig. 1F). As expected, the MYC^high^ group was also enriched with gene sets of MYC targets, the cell cycle and cell growth-related signatures. We next investigated whether the MYC^high^ status of cells has a discernible impact on the expression of genes that encode mitochondrial proteins. For GSEA analysis, we chose a gene set representing MYC-regulated nuclear genes that encode mitochondrial proteins (collated by Morrish & Hockenbery ^32^) and compared the representation of this signature between the MYC^high^ (+++, n=4) and MYC^low^ (0, n=3) TNBC cells (Fig. 1E). The analysis revealed a significant enrichment of MYC-regulated mitochondrial targets in the MYC^high^ group (Fig. 1G).

### MYC^high^ cancer cells show increased mitochondrial respiration

To further explore the association between MYC^high^ status and enrichment of mitochondrial respiration gene sets, we first compiled a list of 26 mitochondrial respiration relevant gene sets identified from various data repositories (MitoCarta3.0 ^33^; MSigDB ^34^, Reactome ^35^). Next, we sourced RNA-seq data for cell lines from Cancer Cell Line Encyclopedia (CCLE; Broad Institute, 1009 cell lines) and for TNBC samples from The Cancer Genome Atlas Program (TCGA; NIH, 105 samples) and, subsequently, computed with the ssGSEA method the enrichment scores (ES) both for the MYC V1V2 signature and separately for all 26 mitochondrial respiration gene sets. The Pearson correlation analysis revealed a significant positive correlation between the MYC^high^ status and 24 out of 26 mitochondrial respiration gene sets in the CCLE data. In the TCGA patient data, MYC^high^ status positively correlated with 21 mitochondrial respiration gene sets out of 26. The most significantly enriched gene sets included OXPHOS, OXPHOS assembly factors, complex I and IV as well as ETC-related gene sets (Fig. 2A, Supplementary Fig. 2A,B). In addition, several other mitochondrial function-related gene sets, such as mitochondrial protein import, translation and ion transport gene set were enriched in the group of MYC^high^ TNBC cells (Supplementary Fig. 2C). Moreover, we also evaluated the association between MYC V1V2 expression and mitochondrial respiration-related gene sets using single cell RNA sequencing (scRNA-seq) data of 10 patients with TNBC (Fig. 2B-D, Supplementary Fig. 2D) and spatial transcriptomics data of 22 patients with TNBC (Fig. 2E, F, Supplementary Fig 2E, F). ssGSVA analysis of scRNA-seq and spatial sequencing revealed high MYC V1V2 expression in TNBC cancer cells and TNBC-enriched spots compared to normal tissue, respectively (Supplementary Fig 2E, D). scRNA-seq analysis showed a strong positive correlation between MYC V1V2 signature and mitochondrial respiration-related signatures in the TNBC cancer cells (Fig. 2D, F). The spatial transcriptomics analysis further confirmed that cancer cell-containing spots with the highest MYC expression also displayed the highest expression of mitochondrial respiration-related signatures (Fig 2F). Supplementary Fig. 3A shows validation of MYC V1V2 signature regarding mitochondrial function-related gene sets. Together, these data reveal a striking association between the MYC^high^ status and transcriptional changes that indicate increased mitochondrial respiration in diverse cancer cell lines and TNBC patient samples.

**Fig. 2.**
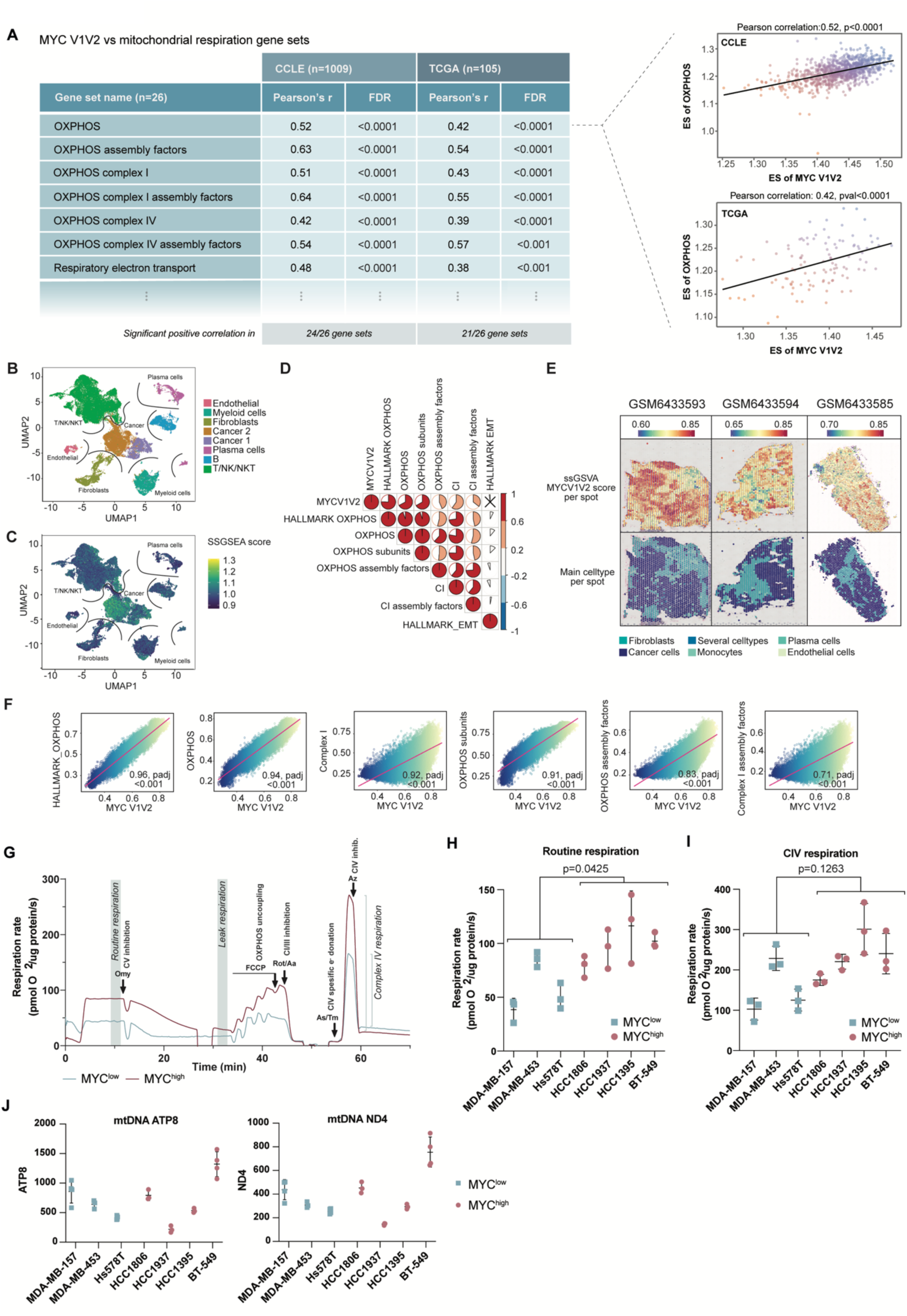
Mitochondrial respiration is increased in MYC^high^ TNBCs. **A** Pearson correlation of MYC V1V2 and OXPHOS enrichment scores (ES) in CCLE and TCGA (TNBC) RNAseq data. Pearson correlation coefficient values, r, and corresponding FDR values for Pearson correlations between ES of several mitochondrial respiration gene sets and ES of MYC V1V2. ESs were calculated from CCLE RNAseq data and TCGA TNBC sample RNAseq data. **B** UMAP of single cell sequencing data with main cell types. **C** UMAP of single cell sequencing data with MYC expression calculated with ssGSVA. **D** Spearman correlation between expression scores of MYC V1V2 and several mitochondrial respiration-related gene sets in scRNAseq data. Only cancer cells were included in the analysis. **E** Spatial sequencing data showing MYC V1V2 expression scores per spot which was defined with using ssGSVA (top images). The images on the bottom show the main cell types per cluster which were defined using seurat prediction algorithm. Spots with a confidentiality score above 0.5 per cell type were considered positive for that specific cell type. If a spot had multiple cell types with a confidentiality score below 0.5, it was considered to have a mixed-cell composition. **F** Spearman correlation between the expression scores of MYC V1V2 and different mitochondrial respiration-related gene sets per spot calculated using ssGSVA. Only spots defined as “cancer” were included into the analysis. **G** Oroboros respiration trace graph. Example respiration trace curves for MYC^high^ (+++) HCC1395 cell line and MYC^low^ (0) MDA-MB-453 cell line measured by Oroboros. Omy = Oligomycin, FCCP = Carbonyl cyanide p-trifluoro-methoxyphenyl-hydrazone, Rot/Aa = Rotenone/Antimycin A, As/Tm = Ascorbate/ N,N,N′,N′-tetramethyl-p-phenylenediamine, Az = Azide. **H** Routine respiration rates measured by Oroboros. Statistical significance: two-tailed unpaired t-test. **I** Complex IV respiration rates measured by Oroboros. Statistical significance: two-tailed unpaired t-test. **J** Mitochondrial DNA (mtDNA) content with mitochondrial target genes ATP8 and ND4 using qRT-PCR assay. Statistical significance: two-tailed unpaired t-test (ns).

To determine whether high MYC activity couples with elevated mitochondrial respiration, we measured the rate of mitochondrial respiration in the MYC^high^ and MYC^low^ groups of TNBC cells using high-resolution respirometry (HRR). The HRR is designed for real-time monitoring of cellular respiration rate and assessment of respiratory complex function via specific substrate-uncoupler-inhibitor titration protocols (example run shown in Fig. 2G). The HRR analyses revealed cell line specific variation within both MYC^high^ and MYC^low^ groups for routine respiration, complex IV-specific respiration and leak respiration (dissipative, non-phosphorylating respiration, where protons leak across inner mitochondrial membrane) rates. Notably, the routine respiration rate was statistically higher in the MYC^high^ than in the MYC^low^ TNBC cells and, also, the complex IV-specific respiration and leak respiration rates trended higher in the MYC^high^ cells (Fig. 2H, I, Supplementary Fig. 2G). Earlier studies have shown in lymphoma cells with conditional MYC that MYC can upregulate respiration in association with mitochondrial biogenesis ^36^. However, in TNBC cells the mitochondrial respiration rate did not appear to correlate with mitochondrial mass in either the MYC^high^ or MYC^low^ groups (Fig. 2J), although the small number of cell lines makes such a comparison difficult. In conclusion, these data suggest that TNBC and cancer cell lines in general with high MYC activity feature elevated mitochondrial respiration.

### Metformin, a weak mitochondrial complex I inhibitor, suppresses proliferation and primes apoptosis preferentially in MYC^high^ TNBC cells

Metformin acts as a weak complex I inhibitor in cell cultures at high concentrations ^37,38^. In accordance, addition of metformin in the millimolar range efficiently inhibited complex I-dependent respiration, with no difference observed between the MYC^low^ and MYC^high^ TNBC cells (Fig. 3A). To explore whether complex I inhibition by metformin has different biological effects in MYC^high^ and MYC^low^ groups of TNBC cells, we defined the proliferation and apoptotic status of metformin treated MYC^high^ (+++) and MYC^low^ (0) TNBC cell lines stratified according to the MYC V1V2 signature. Metformin inhibited proliferation in both MYC^high^ and MYC^low^ groups of cells (Fig 3B). A trend for stronger inhibition of proliferation was observed in the MYC^high^ group, although the difference between MYC^high^ and MYC^low^ cells within each treatment group was not statistically significant. We also measured the level of trimethylation of histone 3 at lysine 9 (H3K9me3), a post-translational modification involved in the formation of transcriptionally silent heterochromatin in senescent cells ^39^. Metformin treatment strongly and selectively induced H3K9me3 marks in the MYC^high^ TNBC cells (Fig. 3C,D). Nevertheless, even though metformin inhibited the proliferation of MYC^high^ cells in short-term (Fig. 3B) and long-term (Supplementary Fig. 4A) assays as well as promoted the senescence-associated H3K9me3 marks (Fig. 3C) and senescence-associated beta-galactosidase activity (Supplementary Fig. 4B, C), the inhibitory effect of metformin on proliferation was reversible (Fig. 3E,F, Supplementary Fig. 4A). Continuous administration of metformin for two weeks inhibited cell proliferation, but proliferation ensued upon withdrawal of metformin (Fig. 3E,F, Supplementary Fig. 4A,D). Our results align with recent findings suggesting that cells can exhibit senescence-associated markers without undergoing the irreversible cell cycle arrest typically defined as senescence ^40^. For instance, studies have shown that the staining intensity of multiple senescence biomarkers is graded rather than binary, reflecting the duration of cell-cycle withdrawal ^41^.

**Fig. 3.**
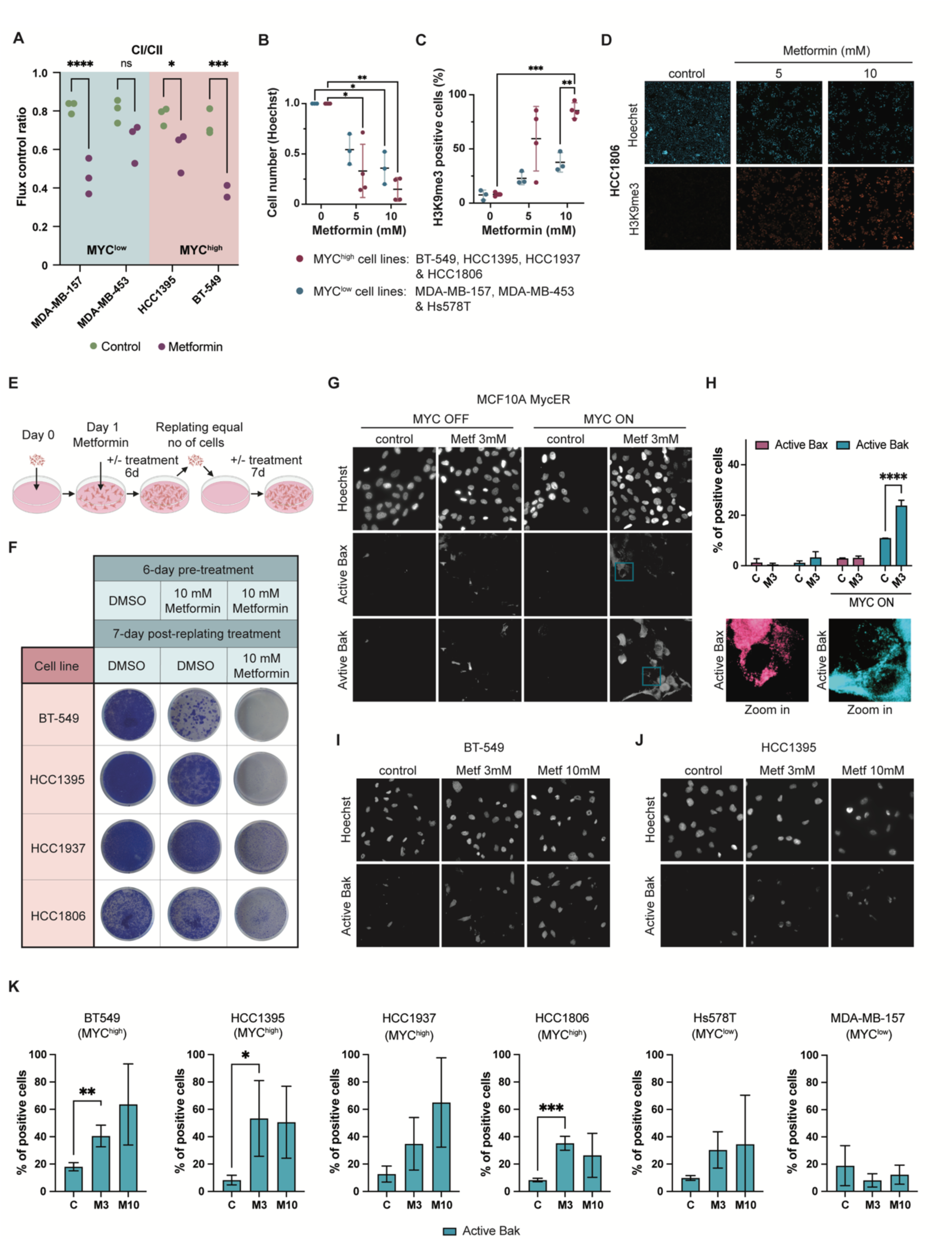
Metformin inhibits proliferation, promotes senescence-associated features, and induces BAK activation in MYC^high^ TNBC cells. **A** Metformin-induced inhibition of complex I-specific respiration measured by Oroboros. Cells were treated with 3 mM metformin for 24 hours. Statistical significance: two-way ANOVA followed by Šídák’s multiple comparisons test. **B** Fold change of cell number per field of view calculated from Hoechst-stained TNBC cells on multiwell plates. MYC^high^ and MYClow cells were treated for 6 days with metformin (5 mM or 10 mM) or control DMSO. Each dot in the figure represents the mean value of one cell line, data derived from three biological replicates. Statistical significance: two-way ANOVA followed by Dunnett’s multiple comparisons test within MYC^low^ and MYC^high^ groups. **C** Percentages of H3K9me3 positive cells were scored from the same experimental data presented in **B**. Statistical significance: two-way ANOVA followed by Dunnett’s multiple comparisons test within MYC^low^ and MYC^high^ groups and two-way ANOVA followed by Šídák’s multiple comparisons test comparing the effect of each concentration between MYC^low^ and MYC^high^ groups. **D** Representative immunofluorescence images of H3K9me3 and Hoechst-stained cells. **E** Schematic illustration of replating assay for senescence. **F** Replating assay for senescence: Images of coomassie blue-stained cells. **G** Images of MCF10A MycER cells stained with conformation specific N-terminal BAK and N-terminal BAX antibodies. Cells with and without induction of MYC were treated for 72 hours with metformin. **H** Quantitative analysis of the experiments shown in **G**, positive cells calculated per field of view. Images from three biological experiments were scored. Representative images of the stained cells shown in the inset. **I, J** Representative images of TNBC cells IF-stained with conformation specific N-terminal BAK antibody. Cells were treated with metformin treatment for 72 hours. Three biological replicates. **K** Quantitative analysis of the experiments shown in **I** and **J**. Statistical significance: two-tailed t-test.

Activation of MYC primes mammary epithelial and tumor cells to apoptotic cell death through a mechanism that involves conformational activation of the pro-apoptotic BCL-2 family protein BAK ^11^. Conformational activation exposes the N-terminal part of mitochondrially localized BAK, which can be detected with N-terminus directed neoepitope-specific antibodies. To determine whether metformin exerts MYC-selective pro-apoptotic effects, we first treated tamoxifen-inducible MCF10A MycER cells (MycER ^12^) with metformin for 72 hours and immunostained the cells with conformation-specific antibodies for activated N-terminal BAK and also for BAX. Metformin induced apoptotic priming observed as increased BAK activation in MYC-induced MCF10A MycER cells, while BAX did not get activated (Fig. 3G,H). Thus, we next focused on the metformin-induced BAK activation in the MYC^high^ and the MYC^low^ TNBC cells. Metformin induced prominent BAK activation in half of the tested TNBC cell lines, with 40-60% of the cells staining BAK positive at 72h timepoint (Fig. 3I-K). All these cell lines were MYC^high^ cell lines. The BAK response to metformin was less prominent in the two MYC^low^ cell lines. This suggests that MYC sensitizes cells to metformin’s pro-apoptotic effects. Conformational activation of BAK and BAX is associated with apoptotic cell death, and in this study, we have not explored the death-priming action of metformin regarding other types of cell death due to the lack of suitable biomarkers for death priming or death induction.

### MYC^high^ TNBC cells are specifically sensitive to Q site-blocking complex I inhibitors

Next, we asked whether the MYC-associated enhanced mitochondrial respiration generates therapeutic vulnerabilities. For this purpose, we performed a drug screen with FIMM Oncology Collection including 526 investigational and approved oncology drugs for TNBC cell lines stratified as MYC^high^ and MYC^low^ (Fig. 4A, Supplementary Fig. 5A). Drug sensitivity scores (DSS) were obtained for all the MYC^high^ TNBC cell lines and for the following MYC^low^ TNBC cell lines: HCC1143 (+), MDA-MB-231 (+), MDA-MB-468 (+) and Hs587T (0). We used both MYC^low^ (0) and MYC^low^ (+) categories as models for MYC^low^ TNBC due to the limited availability of MYC^low^ (0) TNBC cells in the FIMM HTB Unit where we performed the screen. The analyses revealed 105 compounds with high DSS (>10) for MYC^high^ cell lines. These hits included drugs with previously reported synthetic lethal activity with MYC such as inhibitors for cyclin-dependent kinases (Dinaciclib ^3^), mammalian target of rapamycin complex 1/2 (Sapanisertib ^14^) and polo-like kinase 1 (Volasertib and GSK-461364 ^16^) (Fig. 4B). The drugs which showed the highest MYC selectivity (highest difference in DSS between MYC^high^ and MYC^low^ groups) were BAY 87-2243 (CAS 1227158-85-1) and mubritinib (TAK 165; CAS 366017-09-6) (Fig. 4B).

**Fig. 4.**
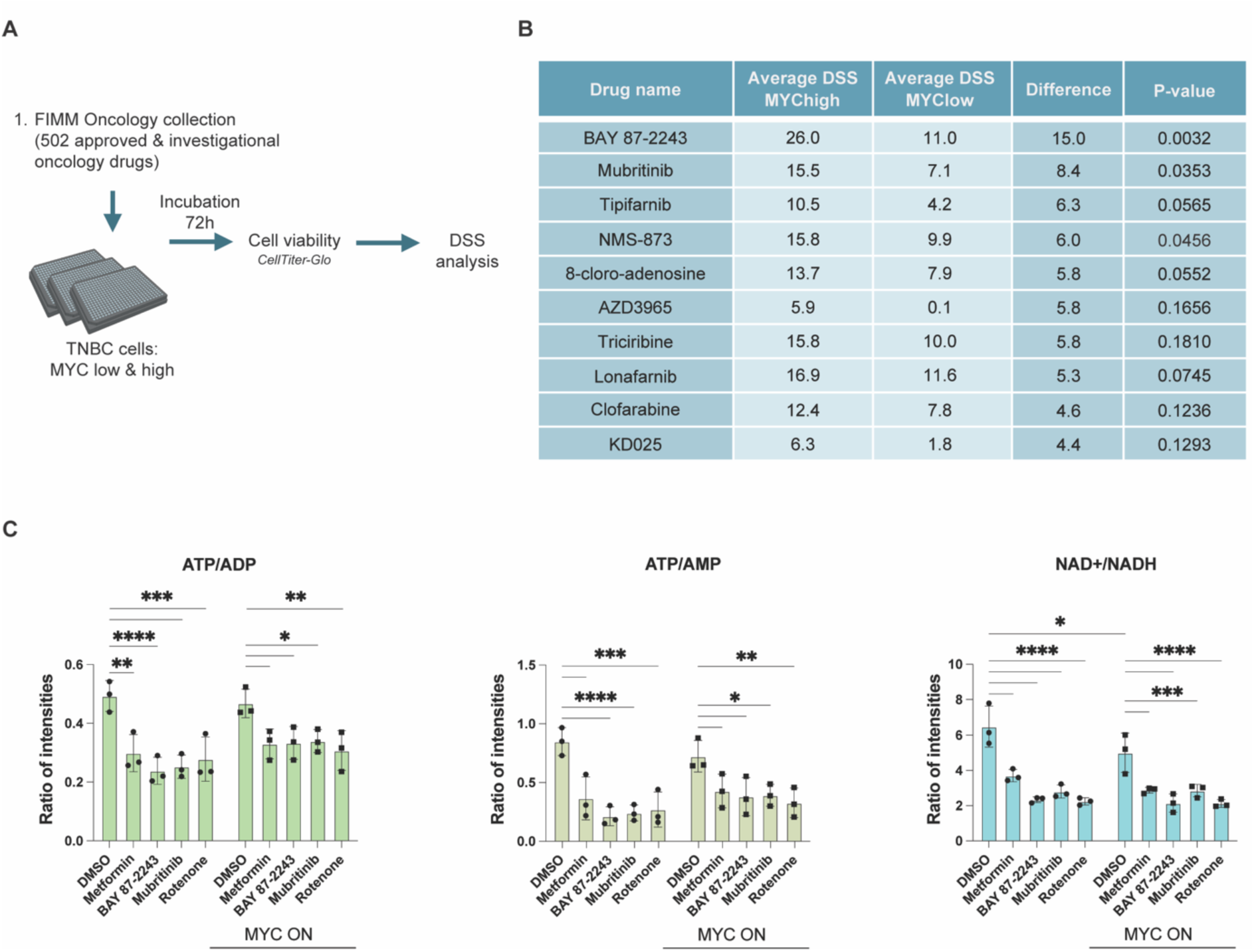
Drug sensitivity screen to identify drugs with selectivity for MYC^high^ TNBC cells. **A** Workflow for the drug sensitivity screen. **B** Top MYC^high^-selective drugs. The difference denotes subtraction of the average drug sensitivity score (DSS) for MYClow TNBC cells from the average DSS for MYC^high^ TNBC cells. The MYC^high^ group included BT-549, HCC1395, HCC1937, HCC1806, HCC1187, MDA-MB436, BT-20 and HCC70 cells and the MYC^low^ group HCC1143, MDA-MB-231, MDA-MB-468 and Hs587T cells. **C** ATP, ADP, AMP, NAD^+^ and NADH intensities (peak areas) from targeted metabolomics assay. MCF10A MycER cells were treated with DMSO, metformin 1 mM, BAY 87-2243 10 nM, mubritinib 10 nM or rotenone 30 nM. MYC was activated with 100 nM 4-OHT. Statistical significance for the MYC OFF *versus* MYC ON controls was calculated using two-tailed unpaired t-test, and for treatments *versus* their respective DMSO controls using two-way ANOVA followed by Dunnett’s multiple comparisons test.

Interestingly, the top two hits, BAY 87-2243 and mubritinib, were originally considered as hypoxia-inducible factor-1 (HIF-1) pathway and receptor tyrosine kinase inhibitors, respectively, and developed for the treatment of cancer. However, later studies have rediscovered BAY 87-2243 and mubritinib as ubiquinone-binding site (Q-site) dependent inhibitors of mitochondrial respiratory complex I ^42–44^. BAY 87-2243 and mubritinib as well as a clinically tested Q-site inhibitor IACS-10759 share an aromatic ring chain topology, a molecular scaffold critical for complex I inhibition ^44^. Our molecular docking analysis of BAY 87-2243, mubritinib and IACS-10759 demonstrated high Q-site binding affinities for all the three compounds (Supplementary Fig. 5B). Furthermore, in MYC^high^ TNBC cells, both BAY 87-2243 and mubritinib induced a prominent AMPK activation, which is a common outcome of complex I inhibition (Supplementary Fig. 5C). These results are consistent with the previous notions that the primary action of BAY 87-2243 and mubritinib is mediated through ubiquinone-binding site (Q-site) dependent inhibition of mitochondrial respiratory complex I.

Complex I a.k.a NADH:ubiquinone oxidoreductase is the first enzyme of the mitochondrial electron transport chain (ETC); it provides entry point for electrons to the respiratory chain by oxidizing NADH to NAD^+^. In the process of mitochondrial respiration, complex I translocates four protons across the inner membrane per molecule of oxidized NADH, helping to build the electrochemical potential difference used to produce ATP. Consistent with this mode of complex I action, we observed that inhibition of complex I with metformin, BAY 87-2243, mubritinib or rotenone in MCF10A MycER cells led to a notable depletion of both NAD^+^ and ATP levels (Fig. 4C). The level of inhibition was similar in these cells with or without MYC activation (Fig. 4C). Therefore, these experiments show that pharmacological inhibition of complex I leads to reductive stress (increased NADH:NAD+ ratio) and loss of ATP, independently of the MYC status.

### MYC enriches the TCA cycle with glutamine-derived carbon

NAD^+^ is an essential coenzyme in multiple pathways, including the TCA cycle, in which dehydrogenase enzymes oxidize their substrates by reducing NAD^+^. Therefore, the inhibition of mitochondrial complex I in MYC-activated cells may lead to disruption in NAD-dependent pathway activity and affect cell survival. To understand how MYC influences the function of the TCA cycle and whether it’s affected by complex I inhibitors, we explored the use of glucose- and glutamine-derived carbons in metabolic ^13^C enrichment analysis.

We first analyzed the enrichment of glucose-derived carbons in glycolysis and TCA cycle intermediates using U-^13^C-labelled glucose tracing in MCF10A MycER cells (Fig. 5A). The pulse isotope tracing showed that MYC induction increased enrichment of ^13^C isotopologues of glycolytic intermediates such as glucose-6P M+6, fructose-6P M+6 and PEP M+3, as measured by both peak areas and overall levels (Fig. 5B, total peak area data table in Supplementary data file 2) as well as their relative abundance to Glucose-6P (Supplementary Fig. 6A). In addition, MYC activation enhanced the enrichment of ^13^C in metabolites of pentose phosphate pathway and glycerol-3-phosphate biosynthesis pathway (Fig. 5B, Supplementary Fig. 6A). Lactate cellular biosynthesis levels, M+3 isotopologue, did not seem to be increased in MYC activated cells (Fig. 5B, Supplementary Fig. 6B). Taken together, the activation of MycER in the MCF10A cells seemed to promote glucose catabolism and pathways branched from glycolytic pathway, consistent with earlier findings of MYC effects on glucose metabolism ^45–47^.

**Fig. 5.**
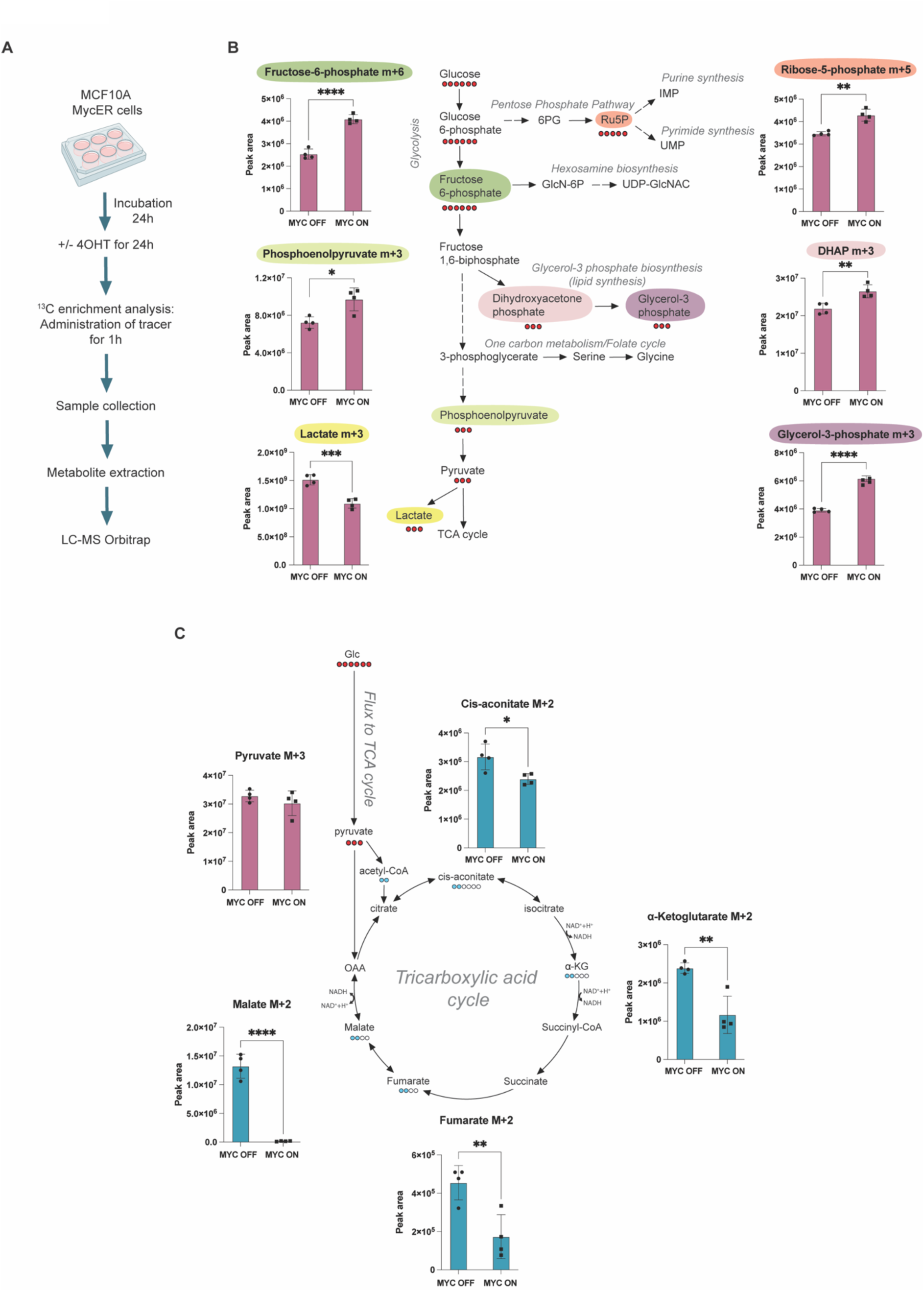
Glucose catabolism and TCA cycle with [U-^13^C]glucose tracer. **A** Workflow for metabolic analysis of MCF10A MycER cells. **B** Glucose-derived ^13^C enrichment in glucose catabolism and branching pathways in nonactivated and MYC-activated MCF10A MycER cells. Statistical significance: two-tailed unpaired t-test. **C** Glucose-derived ^13^C enrichment in TCA cycle. MYC was activated with 100 nM 4-OHT. Statistical significance: two-tailed unpaired t-test. Colored circles indicate the number of ^13^C-labeled carbons in each presented metabolite isotopologue.

Next, we investigated the incorporation of glucose-derived carbons into the TCA cycle in the presence and absence of active MYC (Fig. 5C). We followed the ^13^C enrichment of M+2 isotopologues in the TCA cycle intermediates, which are formed as a result of pyruvate dehydrogenase complex (PDHc) catalysis. We observed that MYC activation reduces the glucose-derived ^13^C M+2 enrichment in α-ketoglutarate, succinate, fumarate and malate (Fig. 5C, Supplementary Fig. 6C-E (time series pulse labeling experiment), all isotopologues shown in Supplementary Fig. 6F), suggesting a possible rewiring of carbon metabolism in the TCA cycle. To further investigate this, we explored the contribution of other carbon sources to the TCA cycle upon MYC activation.

Glutamine serves also as a carbon source for the TCA cycle, initially hydrolyzed by glutaminase to produce glutamate, subsequently converted into α-ketoglutarate before entering the TCA cycle (Fig. 6A). To follow glutamine carbon enrichment in the cells with or without active MYC, we used ^13^C-labelled glutamine, finding that MYC activation significantly increases the glutamate M+5 levels and its relative abundance to glutamine M+5 (Fig 6B, Supplementary Fig 7A,B). A similar increase was observed in the downstream TCA cycle intermediates, with the M+5 isotopologue of α-ketoglutarate and the M+4 isotopologues of succinate, fumarate, and malate, as expected from the first clockwise turn of the TCA cycle (Fig. 6C, all isotopologues shown in Supplementary Fig. 7C). These findings suggest that MYC rewires glutamine pathway to support carbon utilization in the TCA cycle.

**Fig. 6.**
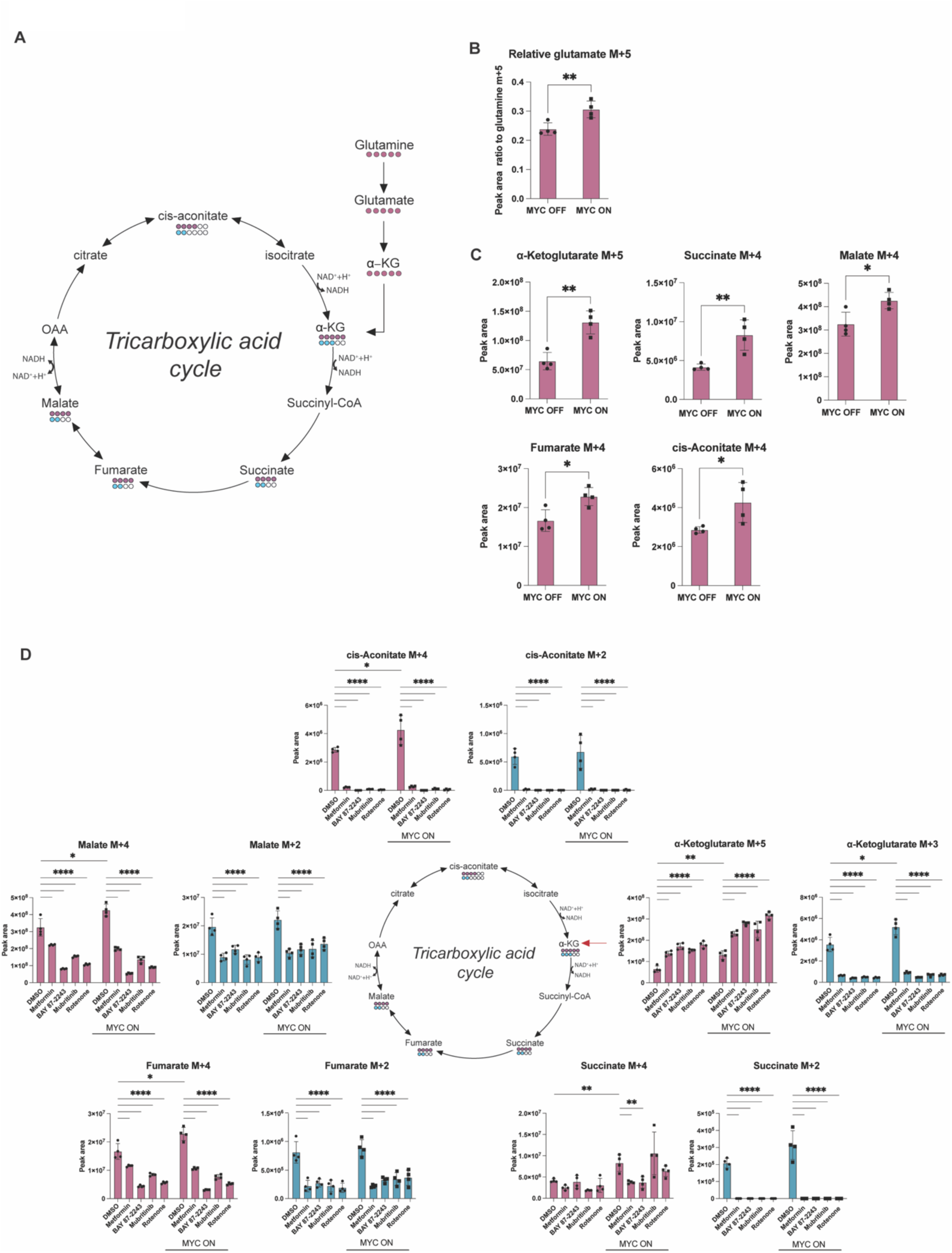
Complex I inhibitors impede MYC-dependent enhanced glutamine carbon incorporation into the TCA cycle. **A** Entry of [U-^13^C]glutamine carbons into the TCA cycle. **B** Relative glutamate M+5 levels to glutamate M+5 levels in MCF10A MycER cells with or without active MYC. Statistical significance: two-tailed unpaired t-test. **C** Glutamine-derived ^13^C enrichment in TCA cycle of MCF10A MycER cells with or without active MYC. Statistical significance: two-tailed unpaired t-test. **D** Glutamine-derived ^13^C enrichment in TCA cycle. MCF10A MycER cells were treated with DMSO, Metformin 1 mM, BAY 87-2243 10 nM, mubritinib 10 nM or rotenone 30 nM. The DMSO controls for MYC ON and OFF are from the same experiment with results shown in **C**. **D** additionally shows the effect of complex I inhibitors in the presence or absence of MYC activation. Statistical significance between DMSO control and inhibitors was calculated using two-way ANOVA followed by Dunnett’s multiple comparisons test. Colored circles indicate the number of ^13^C-labeled carbons in each presented metabolite isotopologue.

Next, we asked whether complex I inhibition disrupts glutamine-derived carbon enrichment in the TCA cycle of MYC-activated cells. In the stable isotope ^13^C-glutamine tracing we found that the complex I inhibitors increased α-ketoglutarate M+5 levels (Figure 6D, Supplementary Fig. 7C) and decreased TCA cycle intermediate M+4 and M+2 isotopologues (Fig. 6C). This indicates that complex I inhibitors can efficiently impede the incorporation of glutamine-derived carbons into the TCA cycle intermediates, regardless of MYC status.

### MYC reprogrammed TCA cycle exposes synthetic lethal opportunities for complex I inhibitors

Finally, we explored the potency of complex I inhibition to induce death in cells with high MYC activity. We first analyzed the effects of complex I inhibition alone or in glutamine-deprived conditions in MCF10A MycER cells using the CellTox Green cell death assay. We observed a prominent MYC-dependent induction of cell death in glutamine-depleted conditions or in the presence of the glutaminase-inhibitor CB-839 (Fig. 7A,B).

**Fig. 7.**
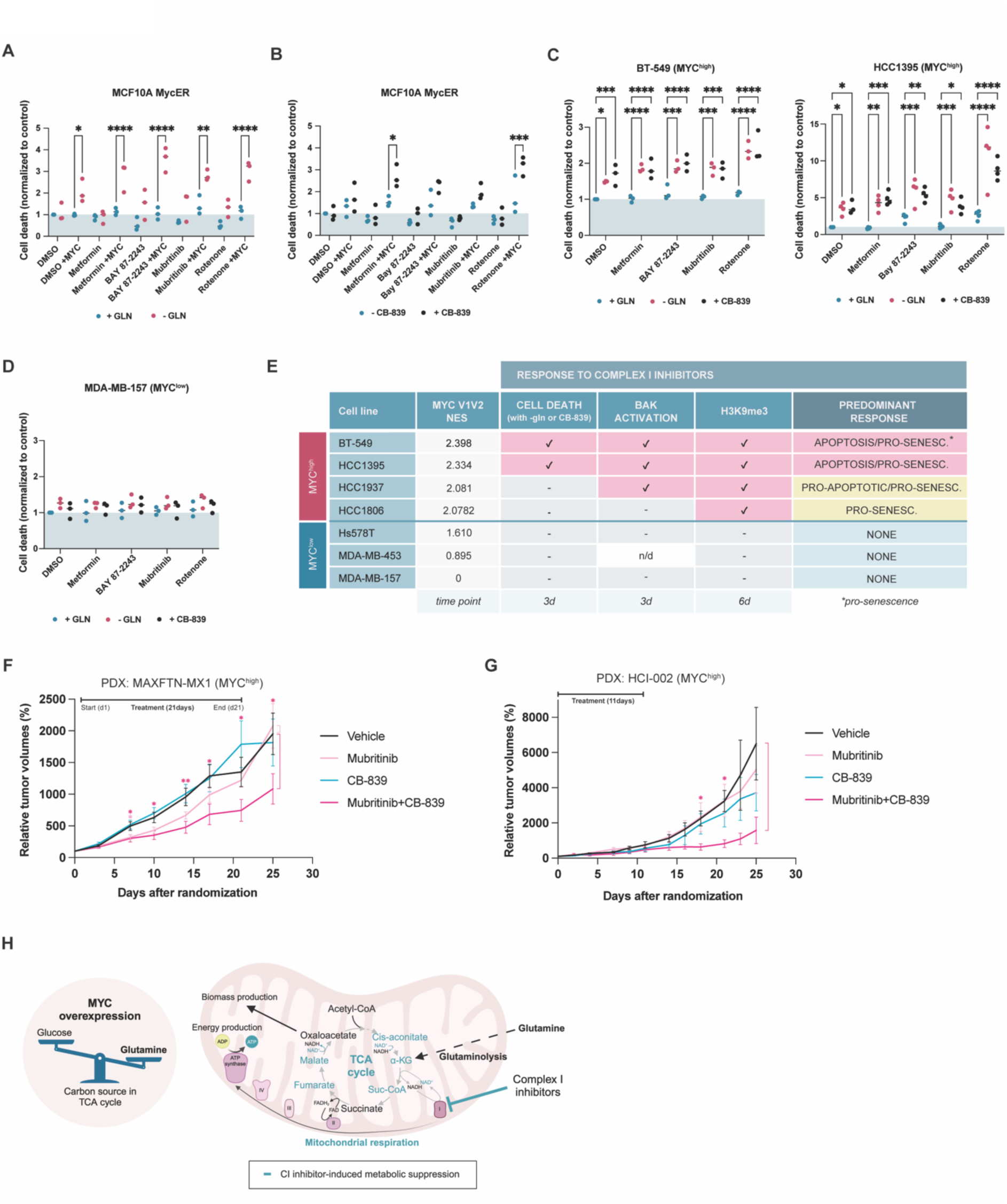
Complex I inhibition with glutamine deprivation preferentially kills MYC^high^ breast cancer cells, inhibiting tumor growth. A,B. CellTox Green cell death assay in MFC10A MycER cells. Cells were treated with DMSO, metformin 1 mM, BAY 87-2243 10 nM, mubritinib 10 nM or rotenone 30 nM with or without active MYC. In **A** cells were cultured in normal growth medium +/-GLN. In **B** +/-glutaminase inhibitor CB-839. Cell death was analyzed at 48 h (A) or 72 h (B) with IncuCyte S3 Live-Cell Analysis Instrument (Sartorius). Statistical significance: two-way ANOVA followed by Šídák’s multiple comparisons test. **C**,**D** CellTox Green assay in MYC^high^ and MYC^low^ cells. Cells were treated as in a-b. Cell death was analyzed at 72h with IncuCyte. Statistics: two-way ANOVA followed by Dunnett’s multiple comparisons test. **E** Summary: Responses of MYC^high^ and MYC^low^ cells to complex I inhibition. Cell death: >50% of the tested combinations induced significant increase in cell death (Fig. 7, C and D). Combinations: complex I inhibitor +/-glutamine deprivation or complex I inhibitor +/-CB-839. BAK activation: >40% of the metformin-treated cells active BAK positive (Fig. 3J). H3K9me3 positive: >75% of metformin-treated cells express H3K9me3 marker (Fig. 3B). **F** Relative tumor volumes of MYC^high^ MAXFTN-MX1 PDXs (n=8/group). Statistical significance: one-tailed Mann-Whitney test. **G** Relative tumor volumes of MYC^high^ HCI-002 PDXs. N=6 (vehicle), n=6 (Mubritinib), n=3 (CB-839) and n=4 (Mubritinib+CB-839). Statistical significance: unpaired t-test with Welch’s correction. **H** A model for the synthetic lethal action of complex I in MYC^high^ breast cancer cells. MYC renders tumor cells dependent on anaplerotic glutamine, which is utilized in TCA cycle. Inhibition of complex I blocks NAD^+^ production, resulting in stagnation of the TCA cycle and inefficient use of the glutamine-derived carbon for anabolic processes, critical for the viability of MYC^high^ cancer cells.

We further tested the lethal effects of complex I inhibition in endogenously MYC^high^ and MYC^low^ TNBC cell lines. The experiments revealed consistently significant induction of cell death in two out of four MYC^high^ cells grown in glutamine-restricted conditions (Fig. 7C, Supplementary Fig. 8A). In the group of MYC^low^ TNBC, two out of three cell lines did not show response to the treatments (Fig. 7D, Supplementary Fig. 8A). The results with endogenously MYC^high^ and MYC^low^ TNBC cell lines are summarized in Fig. 7E, suggesting that complex I inhibitors preferably exert pro-apoptotic and anti-proliferative actions in the MYC^high^ group. The MYC-directed pro-apoptotic effect of complex I inhibitors is closely tied to perturbed glutamine metabolism, which could be one reason to the loss of viability in cells with the MYC^high^ molecular phenotype.

It should be noted that even if these results suggest a MYC-preferred action for complex I inhibitors in a small group of TNBC cells, there was variability in the responses within both MYC^high^ and MYC^low^ TNBC groups. This variability is likely a reflection of the genetic heterogeneity of TNBC cells, which has its own cell line-specific impact on drug responses. Therefore, future studies with a larger panel of cell lines are needed to precisely define the predictive value of the MYC^high^ molecular phenotype for cellular response to complex I inhibitors.

To validate the observed enhanced antitumor activity of complex I inhibition in MYC^high^ tumors *in vivo*, we analyzed a panel of patient-derived xenograft (PDX) models of TNBC from Charles River Laboratories for MYC protein expression (IHC) and MYC V1V2 signature strength (Supplementary Fig. 8B). We chose MAXFTN MX1 as a model of MYC^high^ TNBC for the *in vivo* experiments. As a second MYC^high^ TNBC model, we used HCI-002 PDX (Huntsman Cancer Institute, University of Utah) with earlier defined MYC^high^ molecular phenotype ^3^.

The experiments were performed independently in two different sites (see materials and methods) with slightly modified protocols. Briefly, PDX tumors were implanted subcutaneously (MAXFTN MX1) to NMRI nu/nu mice and allowed to reach about 100 mm^3^ size after which the mice were treated with mubritinib, CB-839 or the mubritinib+CB-839 combination for 21 days and the tumor growth was followed until ethical endpoint. HCI-002 PDX tumors were orthotopically implanted to 4^th^ mammary gland and allowed to reach about 70 mm^3^ after which the tumors were treated for 11 days, and the tumor growth was followed until ethical endpoint.

In both models of MYC^high^ TNBC, mubritinib+CB-839 combination significantly inhibited tumor growth (Fig. 7F,G). On day 25, the relative volumes of mubritinib+CB-839 combination treated tumors were about two (MAXFTN MX1) or five times (HCI-002) smaller than in the control group. Consistent with the analysis of tumor growth, the longest survival time was observed in the combination treated cohorts, yet the differences between groups were not powered (Supplementary Fig. 8C,D). Taken together, it appears that pharmacologic inhibition of complex I together with glutaminase inhibition efficiently suppresses the growth of MYC^high^ tumors. A model to explain the MYC-selective antitumor effect of combined complex I and glutaminase inhibition is presented in Fig. 7H.

## DISCUSSION

Here we demonstrate that MYC V1V2 activity signature, a combination of the two hallmarks MYC target sets, accurately predicts the MYC protein expression status in TNBC cell lines. This signature tool enabled stratification of TNBC cell lines into MYC^high^ and MYC^low^ activity groups, to explore pathways associating with the MYC^high^ phenotype. MYC^high^ cells showed enrichment of cell cycle activity signatures as well as enrichment of DNA repair, cell growth (mTOR), stress and interferon response gene sets. Interestingly, the MYC^high^ signature also indicated association with enhanced oxidative phosphorylation (OXPHOS) and enrichment of MYC-regulated mitochondrial targets ^32^. Consistent with the enriched OXPHOS signatures, MYC^high^ cells showed generally elevated mitochondrial respiration rates. The association between the MYC^high^ and OXPHOS signature was universal, since data analysis of 1009 cell lines in Cancer Cell Line Encyclopedia and 105 TNBC tumor samples in The Cancer Genome Atlas Program demonstrated a significant association between the MYC^high^ signature and OXPHOS- or respiration-related signatures. The association between MYC^high^ status and respiration signatures was also corroborated in single cell and spatial transcriptomics analysis of TNBC tumor samples. These findings universally couple high MYC activity in the cancer cells to persistent upregulation of OXPHOS pathways and enhanced mitochondrial respiration. Earlier findings have presented evidence for association between MYC and OXPHOS in chemotherapy-resistant cancer stem cells of TNBC ^48^ and diffuse large B-cell lymphoma ^49^. High respiratory activity in cancer cells might sound counterintuitive to the concept of Warburg effect that emphasizes the use of non-mitochondrial glycolytic pathways for energy production in tumor cells even if oxygen is present. However, it is well established today that many cancers have elevated mitochondrial activity, as indicated by the enrichment of mitochondrial metabolic signatures or enhanced mitochondrial energy generation. Respiration proficiency has been reported, for example, in diffuse large B cell lymphoma (OXPHOS-DLBCL subtype), melanoma, colorectal cancer, and glioblastoma ^50–53^.

Why would high MYC activity promote mitochondrial respiratory activity? We propose here that the higher respiration rate is part of the MYC’s metabolic program geared to facilitate the anabolic function of the TCA cycle. Mitochondrial respiration produces not only ATP through oxidative phosphorylation but also NAD^+^ from the mitochondrial pools of NADH via complex I activity. The generation of mitochondrial NAD^+^ is in turn indispensable for the function of

TCA cycle due to TCA cycle’s dehydrogenase enzymes, which use NAD^+^ as hydride acceptor ^54,55^. Consistent with the idea that mitochondrially generated NAD^+^ is crucial for sustaining the TCA cycle, we demonstrate here that all tested complex I inhibitors reduced the NAD^+^/NADH ratio and concomitantly the levels of TCA intermediates, exposing a general stagnation of the TCA cycle. Importantly, the TCA cycle is amphibolic, meaning that it has a catalytic function but also an anabolic function crucial for highly proliferative cells such as the cancer cells ^56^. For example, the TCA metabolites are used for the synthesis of amino acids, fatty acids, sterols as well as purines and pyrimidines ^56^. Therefore, coupling of high MYC activity to enhanced respiration benefits cell proliferation in at least two ways: it simultaneously promotes the OXPHOS for production of bioenergy and the use of the TCA cycle for production of biomass.

The metabolic analysis revealed significant differences in the cells’ utilization of glucose- and glutamine-derived carbon in the presence or absence of active MYC. In the presence of active MYC, the glucose-derived carbons were metabolized through glucose catabolism and pathways branched from glycolytic pathway. MYC enriched glucose-derived carbon in the glycolytic pathway and the pentose phosphate pathway, supporting purine and pyrimidine nucleobase synthesis. MYC also enriched glucose-derived carbon in the glycerol-3-phosphate synthesis pathway, which contributes to glycerolipid synthesis. Cellular lactate levels were lower in MYC activated cells, consistent with previous research showing, that MYC enhances expression of monocarboxylate transporters MCT1 and MCT2 exporting lactate from cells and increases excretion of lactate ^17,57^. The decrease in lactate levels could also be due to decreased lactate biosynthesis or shunting of lactate to other pathways. Notably, high MYC activity drastically reduced enrichment of glucose-derived carbon in the TCA cycle. The levels of unlabeled (M+0) TCA cycle intermediates were essentially similar between the cells with and without active MYC with the exception of α-ketoglutarate (M+0), which was increased upon MYC activation. α-ketoglutarate can be derived from glutamine, and further metabolic ^13^C tracer analysis with glutamine-derived carbon showed that MYC significantly enhances the presence of glutamine-derived carbon in all steps of TCA cycle. Together, these findings suggest that in the presence of active MYC, cells prioritize glucose-derived carbon through glucose catabolism and its branched pathways, while relying on glutamine as the primary carbon source for the TCA cycle. This may allow cells to divert glucose-derived carbons into alternative metabolic shunts. Earlier studies have shown that MYC stimulates mitochondrial glutaminolysis, increases the presence of glutamine-derived carbon in the TCA cycle intermediates and renders cells highly sensitive to apoptosis in low glutamine conditions ^17,58,59^.

Here we show that high MYC levels promote switch from the glucose-derived carbon to glutamine-derived carbon as a fuel for the TCA cycle, rendering the TCA cycle mostly glutamine carbon-driven. Therefore, glutamine dependency of MYC^high^ cells should be considered as a major MYC-dependent therapeutic vulnerability.

We demonstrate that the MYC-driven glutamine-fueled TCA cycle can be blocked with different complex I inhibitors and these treatments selectively sensitize cells with high MYC activity to cell death. Even low doses of complex I inhibitors induced substantial cell death in glutamine restricted MYC^high^ TNBC cells. Complex I is a major contributor to mitochondrial NAD^+^ regeneration, which is indispensable for the function of TCA cycle. Therefore, we propose that the lethal action of complex I inhibitors towards the cells with glutamine-fueled TCA cycle arises from deficiency in the oxidizing power of NAD^+^ and consequently, from the impairment of the TCA cycle. The impaired TCA cycle may fail to meet the heightened demand for biosynthesis, anaplerotic use of glutamine and energy production in the MYC^high^ cells with lethal consequences (Fig. 7J). Earlier findings have also highlighted the crucial role for complex I inhibition produced free oxygen radicals (ROS), which, together with the pro-oxidative effects of MYC, could have synthetic lethal effects ^60^. Our results do not refute the role of ROS in mediating TNBC cell death and, hopefully, will inspire future studies to clarify the relative contributions of reductive and oxidative stress in mediating the synthetic lethal action of complex I inhibitors in different cell types with elevated MYC expression.

Several different types of mitochondrial complex I inhibitors have been tested for the treatment of cancer in clinical trials. No results are available for BAY 87-2243 or mubritinib, but the results with structurally related IACS-010759 have raised safety concerns as the treated patients have presented similar spectrum of symptoms observed in patients with mitochondrial diseases caused by deficit OXPHOS ^61^. Therefore, for clinical success in treatment of cancer, the efficacy of safe metformin-like drugs should be improved without simultaneously increasing the risk for adverse effects. Alternatively, biomarkers should be established to identify cancer types sensitive to low doses of ubiquinone-site targeted drugs. Here we show data that could inspire metformin or CI inhibitor combination strategies with glutaminase inhibitors such as CB-839 for the treatment of MYC^high^ cancers. For ubiquinone site-directed complex I inhibitors, we find it encouraging that even low, nanomolar range concentrations of mubritinib induced significant MYC-associated cell death in glutamine-deprived TNBCs. Recent studies have also demonstrated MYC-dependent synthetic lethal activity at nanomolar range of IACS-010759 doses in B-cell lymphoma cells ^8^. With the caution that any comparison of the drug effects between cell culture conditions and *in vivo* is extremely artifactual, the 10 nM dose showing MYC^high^ selective cytotoxicity is about 20-times lower concentrations than the plasma concentration of mubritinib reported to delay development of AML (acute myeloid leukemia) in mouse studies ^42^. Thus, these results may offer a biological rationale for testing low doses of mubritinib ^62^ or other complex I inhibitors in treatment of MYC^high^ cancers or cancer models through dose de-escalation design ^63^. Finally, our mouse studies demonstrate relatively weak (MAXFTN MX1) or no antitumor effect (HCI-002) for mubritinib as a single agent while the combination of mubritinib+CB-839 combination persistently inhibited tumor growth. These results may open new avenues for development of complex I-inhibitor based anti-cancer therapies with persistent efficacy and tolerable adverse effects.

## METHODS

### Cell lines and cell culture

All cell lines were obtained from ATCC. Cell lines were regularly tested for mycoplasma contamination. The cell lines were authenticated by using GenePrint 24 System kit (Promega, B1870) at Institute for Molecular Medicine Finland (FIMM) Genomics unit (HiLIFE infrastructures, University of Helsinki and Biocenter Finland). All cells were cultured at +37°C in a humidified 5% CO_2_ incubator. BT-20 cells were cultured in EMEM medium (Lonza, BE12-125F) supplemented with supplemented with 10 % fetal bovine serum (Serana, S-FBS-SA-015), L-glutamine (Gibco, 25030024), 2 ug/mL insulin (Sigma-Aldrich, I9278) and 100 U/mL penicillin and 100 mg/mL streptomycin (Thermo Fisher Scientific, 15140122). BT-549 were cultured in RPMI 1640 medium (Life Technologies, 31870074) supplemented with 10 % fetal bovine serum (Serana, S-FBS-SA-015), L-glutamine (Gibco, 25030024), 2 ug/mL insulin (Sigma-Aldrich, I9278) and 100 U/mL penicillin and 100 mg/mL streptomycin (Thermo Fisher Scientific, 15140122). HCC1143, HCC1187, HCC1395, HCC1806, HCC1937, HCC38, HCC70, MDA-MB-157, MDA-MB-231, MDA-MB-436, MDA-MB-453 and MDA-MB-468 were cultured in RPMI 1640 medium (Life Technologies, 31870074) supplemented with 10 % fetal bovine serum (Serana, S-FBS-SA-015), L-glutamine (Gibco, 25030024) and 100 U/mL penicillin and 100 mg/mL streptomycin (Thermo Fisher Scientific, 15140122). Hs578T were cultured in DMEM medium (Sigma Aldrich, D6546) supplemented as RPMI medium. For mRNA sequencing MDA-MB-157, MDA-MB-231, MDA-MB-436, MDA-MB-453 and

MDA-MB-468 cells were cultured in Leibovitz’s L-15 medium (ATCC, 30-2008) supplemented 10% fetal bovine serum (Biowest) and 100 U/mL penicillin and 100 mg/mL streptomycin (Thermo Fisher Scientific, 15140122), and MDA-MB-436 cell line was in addition supplemented with 10 µg/mL insulin (Sigma-Aldrich, I9278) and 16 µg/mL glutathione (Sigma).

Mammary epithelial MCF10A cells (obtained from ATCC) were cultured in DMEM/F-12 without phenol red medium (Thermo Fisher Scientific, 21041025), supplemented with 1% horse serum (Thermo Fisher Scientific, 26050088), 10 ng/mL human epidermal growth factor (Sigma-Aldrich, E9644), 0.5 mg/mL hydrocortisone (Sigma-Aldrich, H4001), Cholera Toxin (100 ng/mL (Sigma-Aldrich, C-8052), 5 mg/mL insulin (Sigma-Aldrich, I9278) and 100 U/mL penicillin and 100 mg/mL streptomycin (Thermo Fisher Scientific, 15140122).

MCF10A MycER cells containing pBabe-puro MycER^TM^ construct encoding a 4-hydroxytamoxifen (4-OHT) inducible MYC fusion protein MycER, have been previously described (12).

### Treatments

Cells were treated with the following reagents in the experiments. (Z)-4-Hydroxytamoxifen (4-OHT) (Sigma Aldrich, H7904) was used to activate MycER. Dimethyl Sulfoxide (DMSO) (Sigma Aldrich, D8418) was the vehicle control. The inhibitors used were metformin (MedchemExpress, HY-17471A), BAY 87-2243 (MedChemExpress, HY-15836), mubritinib (MedChemExpress, HY-13501), rotenone (MedChemExpress, HY-B1756) and CB-839 (MedChemExpress, HY-12248).

### *MYC* amplification status

*MYC* amplification status was obtained from Cancer Cell Line Encyclopedia (CCLE) by Broad Institute ^64^.

### RNA sequencing

Total RNA was extracted using RNeasy Mini Kit (74104, Qiagen) according to the manufacturer’s instructions. ScriptSeq Complete Gold Kit (Human / Mouse / Rat) (BG1224, Epicentre) was used for RNA-sequencing library preparation from 1000ng total RNA. Ribosomal RNA was first removed from the total RNA using Ribo-Zero Gold rRNA Removal reagents (RZG1224/MRZ11124C, Epicentre), after which the ScriptSeq v2 RNA-Seq Library Preparation Kit (SSV21124, Epicentre) was used to make the Rna-seq library from the Ribo-Zero treated RNA. The rRNA-depleted RNA was fragmented chemically, and the libraries were prepared according to the manufacturer’s instructions. Finally, the libraries were assessed using Agilent High Sensitivity D1000 ScreenTape Assay with 4200 TapeStation System. Samples were sequenced with the Illumina NextSeq 500 instrument using 75bp paired-end reads with an average sequencing depth of 19M reads/sample

### Differential gene expression and gene set enrichment analysis

Differential expression analysis of genes was performed for the RNA-seq data in all the cell lines against MD-MB-157 reference cell line using edgeR BioConductor package ^65^. Gene Set Enrichment Analysis (GSEA; ^66^) and Correlated Adjusted MEan RAnk gene set test (CAMERA; ^67^) were used for functional enrichment analyses based on the differential expression results. Gene sets for functional analysis were chosen from the MSigDb database ^31,34^. Specifically, the hallmark gene sets, gene ontology gene sets, oncogenic signature gene sets and MYC-related gene sets were included in this analysis. Additionally, Chandriani and Goga MYC signatures ^13,25^ were also included in the analysis to carry out a thorough analysis of known MYC signatures in literature. In total, 10,453 gene sets were included in the functional analyses. For GSEA analysis, the ranking metric used is as follows: - sign(log2FC) * log10(P). R package pheatmap was used to perform the hierarchical clustering with Euclidean distance.

### MYC protein level definition

To assess the relative levels of MYC protein expression in TNBC cell lines, the cells were plated on poly-D-lysine coated 96-well plates (354640, Corning BioCoat) and allowed to settle for 53 hours. Cell line-specific optimized seeding densities were scored from previously established growth curves measured with IncuCyte S3 (Sartorius) live-cell analysis, to ensure that the cultures remained sub-confluent over the whole 53-hour period of the experiment. The cells were fixed with 3.7% PFA for 15 minutes, washed three times with PBS and permeabilized with 0.1% Triton X-100 in PBS (PBST). For blocking, we used 5% bovine serum albumin in PBST for 1 hour at RT. The cells were incubated overnight at +4°C with 1:500 MYC antibody (ab32072, Abcam) diluted in 5% BSA-PBST. Thereafter, the cells were washed with blocking Buffer (5% BSA in PBS) and the secondary antibody (Goat anti-Rabbit IgG; 84541, Dylight) diluted 1:500 in blocking buffer was added and the cells were incubated for 1 hour at RT in the dark. The cells were washed with 5% BSA in PBS. The nuclei were stained with 1:10000 Hoechst (H33258, Sigma) for 10 minutes. Nuclear MYC staining was analyzed using Thermo Scientific Cellomics ArrayScan HCS analysis system. To obtain the protein expression score (PES), the mean values of MYC protein expression for each TNBC cell line were compared to MDA-MB-157 cell line and the results were expressed as fold-change. Spearman correlation analysis was computed for the MYC protein expression scores (PES) and the normalized enrichment scores (NES) obtained by GSEA.

### Single cell RNA and spatial transcriptomics

The data used for spatial and single cell sequencing analysis was downloaded from GEO with accession numbers: GSE210616, GSE176078 and GSE161529 ^68–70^. The analysis was performed with R version 4.4.2, R-studio and Seurat 5.1.0 and followed the guidelines of Seurat-package ^71–75^.

For the single cell RNA sequencing analysis, a set of 15 TNBC samples was used. Patients with prior neoadjuvant treatment were excluded from the single cell analysis. From the single cell data, low-quality cells were removed by filtering out cells with less than 300 counts and 500 features, and by removing cells with more than 20% of mitochondrial genes. Duplets were excluded by removing cells with over 5000 counts. In the scRNA analysis, the total set consisted of 52893 cells, that passed the quality control, and of which 13316 were classified as epithelial cells. Single cell data was normalized with SCTransform ^76,77^, integrated with Harmony, and individual clusters were calculated with resolution 0.3 and identified by using conventional markers for each cell type presented in the Supplementary Fig. 9A. Optimal number of clusters was defined by clustree algorithm ^78^.

A total of 56567 spots from 43 slides derived from 22 patients were selected for the spatial analysis. The data was normalized with SCTransform. Tumor-containing regions were identified from the spatial transcriptomics data using three distinct methods with cells from scRNA-seq samples MH0135 and CID44991 serving as references. The accuracy of the identified regions was validated by calculating the purity score using the ESTIMATE ^79^ (Supplementary Fig. 9B). Eventually spots with over 0.5 prediction scores for cancer cells in single cell reference mapping were defined as cancer tissue and selected for further analysis.

The individual gene sets used for the study were downloaded either form MitoCarta3.0 ^33^ or from MSigDB ^66^. The gene set expression was calculated with ssGSVA. Correlations were calculated using Spearman correlation.

### High-Resolution Respirometry

High-resolution respirometry was performed as described ^80^. Briefly, cells were harvested by trypsinization and resuspended in respiration medium MiR05 (0.5 mM EGTA, 3 mM magnesium chloride, 60 mM K-lactobionate, 20 mM taurine, 10 mM potassium dihydrogen phosphate, 20 mM HEPES, 110 mM sucrose and 1 g/L fatty acid free BSA (pH 7.1)). Specific oxygen consumption rates (expressed as pmol O2/s/mg of protein) were determined in non-permeabilized or digitonin-permeabilized cells in a high-resolution respirometer (Oxygraph-2k; Oroboros Instruments, Innsbruck, Austria) following specific substrate-uncoupler-inhibitor titration protocols. The non-permeabilized protocol assessed routine respiration, leak respiration (oligomycin) and electron transfer capacity (carbonylcyanide p-trifluoromethoxyphenyl-hydrazone (FCCP)). Residual respiration was measured by addition of rotenone (0.5 μM) and antimycin A (2.5 μM). Complex IV activity was obtained by adding ascorbate (2 mM) and TMPD (N,N,N,N-Tetramethyl-p-phenylenediaminedihydrochloride) (0.5 mM) followed by azide (10 mM). The permeabilized protocol was initiated by adding digitonin (5 µg/mL). Leak respiration was evaluated by adding malate (1 mM), pyruvate (5 mM) and glutamate (10 mM). For complex I evaluation, ADP+Mg2+ (20 mM) was added. Subsequently, succinate (10 mM) was added for complex II evaluation. Next, all the steps from the non-permeabilized protocol were performed starting at the addition oligomycin. Mean values of oxygen flux at specific respiratory states were determined using DatLab 5 (Oroboros Instruments GmbH, Innsbruck, Austria). All experiments were performed in triplicates per cell line.

### Mitochondrial DNA extraction and qRT-PCR

Mitochondrial DNA (mtDNA) extraction and quantitative real-time PCR assay were performed according to previous studies ^81^. In short, DNA was extracted and purified from cultured cells using QIAamp DNA Mini Kit (Qiagen, 51304) according to the QIAamp DNA Mini Handbook. The concentration of the samples was measured using Qubit dsDNA Quantification Assay Kit (Invitrogen, Q32850) and Qubit 4 Fluorometer. A SYBR green dye-based qPCR (1708882; Bio-Rad) of target genes were performed in the Bio-Rad CFX machine (Bio-Rad). Every sample was run in triplicate. The PCR program used was: 5 min at 95°C initial denaturation, followed by 40 cycles of 10 s of denaturation at 95°C, 30 s of primer annealing at 60°C and 10 s at 72°C of extension. The primers have been listed in Supplementary table 1. MtDNA content was calculated by the comparative CT method, normalizing it to nuclear DNA with geometric mean of two nuclear reference genes CIAO1 and SNW1 followingly ^82,83^:

Relative mitochondrial DNA content= 2*2^-[CT(target)-√(CT(CIAO1)*CT(SNW1))]

### Automated Immunofluorescent imaging of Hoechst and H3K9me3

Immunofluorescence staining was performed to calculate the cell numbers and determine the proportion of H3K9me3-positive cells after selected treatments. Cells were seeded onto Perkin Elmer 96-well View imaging plates. After 24 hours, the wells were inspected under a light microscope to ensure even distribution of the cells. The culture medium was then changed, and the cells were treated with metformin for 6 days. Dimethyl sulfoxide (DMSO) was used as a vehicle control and added to the metformin treatments as well, in low concentration of 0.01%. Cells were washed two times with PBS. Cells were fixed using 4% paraformaldehyde (PFA) solution in PBS for 15 minutes. Then, cells were washed twice with PBS and permeabilized with 0.1% Triton X-100-PBS for 6 minutes, followed by washing. Non-specific binding sites were blocked using 1% BSA-PBS for 1 hour. Cells were incubated with primary antibody H3K9me3 (Abcam, ab8898) in dilution of 1:500 in 1% BSA-PBS overnight in +4°C on a rocker. Cells were washed 3 times and incubated at room temperature for 1 hour with Alexa Fluor 546 conjugated secondary antibody (Life technologies, A11010) in 1:300 dilution in 1% BSA-PBS followed by two washes of the cells. Nuclei were counterstained with 1 mg/mL Hoechst 33258 for 4 minutes. As a last step, 200 uL of phosphate-buffered saline was added to the wells and a sticker was put on top to protect wells. Experiments were performed with technical and biological triplicates.

Plates were imaged using automated-imaging imaging instrument, CellInsight CX5 High-Content Screening (HSC) Platform and later with Molecular Devices ImageXpress Pico Automated Cell Imaging System due to CellInsight being removed from use. Images were taken with 10x objective. Thresholds for imaging were set to image 1000 cells or 36 images or 3 sequential images with <2 cells with CellInsight and 4 images from the center of the well with ImageXpress Pico. Immunofluorescence images of H3K9me3 were quantified using Cellomics HSC Studio 4.0 Cell Analysis Software and Target Activation analysis with the CellInsight images. IN Carta Image Analysis software and 2-channel marker expression analysis was used for images from ImageXpress Pico. Number of the cells were counted using nuclei stain. Percentages of H3K9me3 positive cells were counted according to the intensity of H3K9me3 stain. Intensity threshold was chosen for each cell line by assessing both control wells and treated wells.

### Drug withdrawal and time-lapse video microscopy

Cells were seeded on 96-well plates. 24 hours after seeding cells were treated with metformin and dimethyl sulfoxide was used as a vehicle control and added to the metformin treatments as well, in low concentration of 0.01%. Plates were incubated under 5% CO_2_ and kept at +37°C in a IncuCyte S3 Live-Cell Analysis Instrument (Sartorius) for 15 days. Culture medium was changed every 3 days. Video recordings were taken using 10x objective and by recording 5 frames per well every 2 hours. Analysis and quantification were done using IncuCyte S3 Live-Cell Analysis Instrument (Sartorius). Experiments were performed with technical duplicates and biological triplicates.

### Senescence-associated beta-galactosidase staining

24 hours after seeding cells to 6-well plates, culture media was changed, and cells were treated with metformin or 0.01% dimethyl sulfoxide (DMSO) as a vehicle control for 6 days. Plates were incubated under 5% CO_2_, kept at +37°C and culture medium was changed every 3 days. After six days of metformin treatment, equal number of cells from each condition were replated to new 24-well plates. Treatments were continued for 24 hours. Cells were fixed using 2% PFA, 0.2% glutaraldehyde solution for five minutes and washed with PBS. Cytochemical detection of senescence-associated beta-galactosidase activity was done using chromogenic substrate 5-bromo-4-chloro-3-indolyl β-D-galactopyranoside (X-gal) in a freshly made staining solution (40 mM citric acid/Na phosphate buffer, 5 mM K_4_[Fe(CN)_6_]×3H_2_O, 5 mM K_3_[Fe(CN)_6_], 150 mM sodium chloride, 2 mM magnesium chloride and 1 mg/mL X-gal (Thermo Scientific, R0404) in distilled water) at pH 6.0. Cells were incubated at +37°C overnight. Cells were washed with PBS and 70% glycerol was added on top of cells to prevent drying. Plates were imaged using Eclipse Ts2 (Nikon) color microscope camera. Experiment was performed with biological triplicates.

### Replating assay

Cells were seeded on 6-well plates. 24 hours after the plating, culture media was changed, and cells were treated with metformin for 6 days. Dimethyl sulfoxide was used as a vehicle control and added to the metformin treatments as well, in low concentration of 0.01%. Plates were incubated under 5% CO_2_ and kept at +37°C. Culture medium was changed every 3 days. After 6 days, equal number of cells from each condition were replated to new 24-well plates. Treatments were either withdrawn or continued for 7 days and culture medium was changed every 2 to 3 days. Cells were fixed using 4% PFA solution in PBS for 15 minutes. Cells were washed two times with PBS and stained with Coomassie Brilliant Blue solution (1% Coomassie Brilliant Blue (Sigma), 45% methanol, 9% glacial acetic acid, 45% H20) for 1h, followed by rinsing the plates with PBS and MilliQ H2O. Plates were dried out and scanned. Experiments were performed with biological triplicates.

### Detection of BAK and BAX conformational changes using fluorescent microscopy

Cells were plated on poly-D-lysine coated 8-well chamber slides. 24 hours after seeding cells were treated with metformin for 72 hours. Cells were fixed with 4% PFA solution in PBS for 15 minutes. For immunostainings with conformation-specific antibodies, cells were permeabilized and blocked with 0,08% Saponin + 10% normal goat serum (NGS) in PBS. Primary antibody staining was done using 1:100 dilution of anti-Bak (NT; Upstate, 06-536) and anti-Bax (clone 6A7; Santa Cruz, sc-23959) in 0,06% Saponin + 5% NGS in PBS at +4°C overnight. Cells were washed 3 times and incubated in room temperature for 1 hour with Alexa Fluor 488 or 546 conjugated secondary antibody (Life technologies) in 1:300 dilution in 0,06% Saponin + 5% NGS in PBS. Cells were washed 3 times and nuclei were counterstained using 1 mg/mL Hoechst 33258 for 4 minutes. Cells were mounted using Immu-Mount medium. Immunofluorescent images were aquired in Biomedicum Imaging Unit (University of Helsinki) by using Zeiss Axio Imager.Z2 upright epifluorescence wide-field microscope with 63x objective. Image analysis for positive cells was done by using a pipeline created in CellProfiler (Broad Institute of MIT and Harvard) ^84^.

### CCLE and TCGA RNA-seq data

RNA-seq data for CCLE (Cancer Cell Line Encyclopedia) and breast cancer TCGA (The Cancer Genome Atlas) was downloaded from recount3 ^85^. TNBC patients were selected from Breast Cancer TCGA according to Lehmann et al. 2011 ^86^ and samples with low tumor purity were filtered out. ES (enrichment score) for CCLE and TNBC TCGA data was computed using ssGSEA (single sample GSEA) ^66^.

### Drug Sensitivity Screening

The FIMM FO5A library drug screening was performed essentially as described before in ^87^. Briefly, FIMM Oncology Collection including 526 drugs was used in drug sensitivity screening using the following MYC^high^ and MYC^low^ cell lines. MYC^high^ cell lines: BT-549, HCC1395, HCC1937, HCC1806, HCC38, HCC1187, MDA-MB436, BT-20 and HCC70 and MYC^low^ cell lines: HCC1143, MDA-MB-231, MDA-MB-468 and Hs587T. At the start of the screen, the drugs were administered on 384-well culture plates using acoustic liquid handler. The drugs were administered in five increasing concentrations in 10-fold dilution, covering concentration range of 10,000-fold (e.g. 1-10,000 nM), each concentration as a singlets. The drug concentration ranges have been defined as described in ^88^. Essentially, the drug concentration ranges have been adjusted based on drug responses for primary samples data. The cells were plated by a dispenser and incubated for 72h in +37°C in a humidified 5% CO2 incubator. After the incubation period the cell viability was measured using CellTiter-Glo (Promega). The drug sensitivity scores (DSS,^89^) were determined based on the CellTiter-Glo measurement. The quality control and analysis of the dose-response curves was carried out using FIMM in-house Breeze web-tool ^90^. Averages of sensitivity scores from MYC^high^ cell lines were compared to MYC^low^ cell lines by subtracting MYC^low^ average values from MYC^high^ average values.

### Western blotting

Cells were lysed from 6-well plates to extract proteins using cold RIPA buffer (50 mM Tris-HCl pH 8, 150 mM NaCl, 1% Triton X-100, 0.5% sodium deoxycholate and 0.1% SDS) supplemented with Pierce Protease Inhibitor Mini Tablets (Thermo Scientific, A32953) and phosphatase inhibitor (Pierce Phosphatase Inhibitor Mini Tablets (Thermo Scientific, A32957). After adding RIPA buffer, the cells were scraped from the plates and incubated 10 minutes on ice. The nuclei were broken by a 25 G x 1 inch needle. The cells were centrifuged for 20 minutes at 16800 RCF at +4°C. The concentration of the samples was measured using BioRad DC Protein assay kit. 20 mg protein was used per sample. 5x loading sample buffer was added to the samples and the samples were denaturized at +95°C heat block for 2 minutes.

The samples were loaded into 4–20% Biorad gradient gels and separated by electrophoresis. The proteins were transferred onto a nitrocellulose membrane by Trans-Blot Turbo Transfer System. Equal loading was confirmed by staining the membranes with Ponceau S solution (P7170-1L, Sigma-Aldrich). Then the membranes were washed with Tris-buffered saline, TBS (10 mM Tris-Cl, pH 7.4; 150 mM NaCl). The binding of antibody to unspecific sites was blocked with blocking buffer (5% m/v non-fat dry milk, TBST (10 mM Tris-Cl, pH 7.4; 150 mM NaCl; 0,05% Tween^TM^20)). Membranes were incubated with primary antibodies (Supplementary Table 1) diluted into blocking buffer on a rocker overnight at +4°C. Washes of the membranes were performed with TBST. The membranes were incubated 1 hour with HRP-conjugated secondary antibodies on a rocker. After washes, ECL Western Blotting Substrate (Thermo Fisher Scientific) was used for detection of HRP activity. The signal was detected by ChemiDoc Imaging System (Bio-Rad).

When a total protein antibody was used after a phosphorylation site-specific antibody, the membrane was stripped in between. The membrane was incubated in stripping buffer (62.5 mM Tris-HCl pH 6.8, 2% SDS, 0.4% β-mercaptoethanol) in +56°C oven for 25 minutes with some agitation. After the incubation the membrane was washed under running water multiple times and then washed in TBST for 5 minutes. The protocol was then continued with the blocking step.

### Targeted LC-MS metabolomics profiling analysis

The cells were plated on 6-well plates with parallel wells reserved for counting the cells. 24 hours after the plating medium was changed and complex I inhibitors and 4-hydroxytamoxifen were added for 24 hours. In tracer experiments 17.5 mM of uniformly labelled [U-^13^C]glucose (Cambridge Isotope Laboratories, CLM-1396-1) or 2 mM of uniformly labelled [U-^13^C]glutamine (Cambridge Isotope Laboratories, 1822-H-PK) in Milli-Q water, was added on cells 1 hour before quenching. In the time series pulse labeling experiment, where the [U-13C]glucose and [U-13C]glutamine labeling experiments were repeated for MYC ON-OFF comparisons with timepoints, the tracers were added 15 minutes, 30 minutes, 1 hour or 2 hours before the quenching. After washing twice with PBS metabolites were extracted from 2D MC10A cells using 400 µL of cold LC-MS grade extraction solvent (Acetonitrile:Methanol:MQ; 40:40:20, Thermo Fisher Scientific) with scraper and subsequently, samples were vortexed for 2 minutes and sonicated for 1 minutes followed by centrifugation at 14000 rpm at +4°C for 5 minutes. After centrifugation supernatants were transferred into polypropylene tubes, placed into Nitrogen gas evaporator and dried samples were suspended with 40 µL of extraction solvent (ACN:MeOH:MQ; 40:40:20) and vortexed for 2 minutes and transferred into HPLC glass auto sampler vials. 2 µL of samples were injected with Thermo Vanquish UHPLC coupled with Q-Exactive Orbitrap quadrupole mass spectrometer equipped with a heated electrospray ionization (H-ESI) source probe (Thermo Fischer Scientific). A SeQuant ZIC-pHILIC (2.1×100 mm, 5-μm particle) column (Merck) was used for chromatographic separation. The gradient elution was carried out with a flow rate of 0.100 mL/minutes with using 20 mM ammonium hydrogen carbonate, adjusted to pH 9.4 with ammonium solution (25%) as mobile phase A and acetonitrile as mobile phase B. The gradient elution was initiated from 20% Mobile phase A and 80% of mobile phase B and maintain till 2 minutes, followed by 20% Mobile phase A gradually increasing up to 80% till 17 minutes, then 80% to 20% Mobile phase A decrease in 17.1 minutes and maintained up to 24 minutes. The column oven and auto-sampler temperatures were set to 40 ± 3°C and 5 ± 3°C, respectively. MS was equipped with a heated electrospray ionization (HESI) source using polarity switching and following setting: resolution of 35,000, the spray voltages: 4250 V for positive and 3250 V for negative mode, the sheath gas: 25 arbitrary units (AU), and the auxiliary gas: 15 AU, sweep gas flow 0, Capillary temperature: 275°C, S-lens RF level: 50.0. Instrument control was operated with the Xcalibur 4.1.31.9 software (Thermo Fischer Scientific). The peak integration was done with the TraceFinder 4.1 software (Thermo Fischer Scientific) using confirmed retention times for metabolites (m+0) standardized with library kit MSMLS-1EA (Merck). ^13^C isotopologues were analyzed with change of m/z (m+1, m+2 etc.). The data quality was carefully monitored throughout the run using pooled QC sample prepared by pooling 5 µL from each suspended samples and interspersed throughout the run as every 10th sample. The metabolite data was checked for peak quality, % RSD and carryover. Metabolomics data was normalized to cell number from parallel wells in the 1-hour timepoint experiments and to total ion count in the time series pulse labeling experiment.

### Computational Modeling Studies

The MOE 2022.02 software suite (Chemical Computing Group Inc., Montreal, Canada) was utilized for the modeling and docking experiments of Complex I protein. Two cryo-electron microscopy (cryo-EM) structures were employed in this study: mitochondrial Complex I from Mus musculus bound to IACS-2858 (PDB ID:7B93) and from Bos taurus (bovine) bound to the phenformin derivative IM1761092 (PDB ID:7R45). The initial structure preparation for protein-ligand complex involves several independent steps: first, hydrogens are added and positioned; tethers are assigned; and finally, energy minimization is performed. During the docking evaluation, dynamically generated conformations of Complex I inhibitors were introduced into the ubiquinone binding site (Q-site) by using the Triangle Matcher algorithm, and these conformations were subsequently evaluated with the London dG scoring function, leveraging the Amber10-Extended Hückel Theory force field. The top 30 ranked poses were then subjected to a refinement process involving energy minimization and re-evaluation using the Generalized-Born Volume Integral/Weighted Surface area (GBVI/WSA dG) scoring function.

### Cell Death experiments

The cells were plated on black, clear bottom 96-well plates for the experiments. 24 hours after the plating medium was changed to fresh medium (in glutamine lacking (-GLN) conditions to medium without glutamine), inhibitors, 4-hydroxytamoxifen and CellTox Green Cytotoxicity reagent (Promega, G8731) at 1/2000 concentration were added. The cells were live-imaged up to 72 hours in IncuCyte S3 Live-Cell Analysis Instrument (Sartorius) placed in +37°C in a humidified 5% CO_2_ incubator and analyzed using the Analysis wizard of the system. Quantification was done using the IncuCyte.

### Immunohistochemistry

The MYC expression of the original breast cancer tumors was defined with IHC staining. Tumor tissues were fixed with 4% PFA and embedded in paraffin and cut into 5 mm slices and deparaffinized. The antigen retrieval was performed in the microwave for 20 minutes by using pH 6 citrate buffer (DAKO). The endogenous peroxidase activity was blocked with 3% H2O2 for 20 minutes. The primary MYC antibody (ab32072, Abcam) was used at the concentration 1:300 and the slides were incubated in +4°C for overnight.

### Patient-derived xenografts

The patient-derived xenograft mouse experiment with MAXFTN-MX1 model was executed by Charles River Laboratories, Germany. Mouse experiments were approved by the German Committee on the Ethics of Animal Experiments of the regional council (Permit Numbers: G-20/163). This study was carried out in strict accordance with the recommendations in the Guide for the Care and Use of Laboratory Animals of the Society of Laboratory Animals (GV SOLAS) in an AAALAC accredited animal facility. Four- to six-week-old female NMRI nu/nu mice (Charles River, Germany) placed under isoflurane anesthesia received tumor implants subcutaneously in the right flank. Animals and tumor implants were monitored daily until the maximum number of implants showed clear signs of beginning solid tumor growth. Tumor volumes were calculated according to the following equation: Tumor Vol (mm^3^) = a (mm) × b^2^ (mm^2^) × 0.5, where “a” is the largest diameter and “b” is the perpendicular diameter of the tumor representing an idealized ellipsoid. At randomization, the volume of growing tumors was initially determined. Animals bearing 50 to 250 mm^3^ tumors, preferably 80 to 200 mm^3^, were distributed into experimental groups, with comparable median and mean tumor volumes. The day of randomization was designated as day 0 of the experiment and dosing was started on day 1. The relative volume of an individual tumor on day X (RTVx) was calculated by dividing the absolute volume (mm^3^) of the respective tumor on day X (Tx) by the absolute volume of the same tumor on the day of randomization, i.e., on day 0 (T0), multiplied by 100, as shown by the following equation:

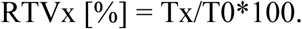

Tumor inhibition on a particular day (T/Cx) was calculated from the median RTV of a test group and the median RTV of a control group multiplied by 100, as shown by the following equation:

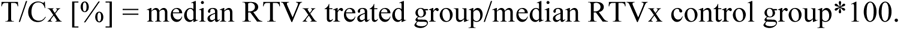

The minimum T/C (%) value recorded for a particular group during an experiment represented the maximum anti-tumor activity for the respective compound (=optimal T/C). Tumor volume and body weight were determined twice per week. Ethical limit for the tumor volumes was 2000 mm^3^. In the ethical protocol, maximum body weight loss was 30% or continued body weight loss of >20% for more than 2 days or rapid body weight loss of >20% within 2 days.

The mouse experiment with HCI-002 model was approved by the National Animal Experiment Board of Finland (Licence number: ESAVI/23309/2022). Four- to six-week-old female NSG (NOD.Cg-*Prkdc^scid^ Il2rg^tm1Wjl^*/SzJ) mice under isoflurane anesthesia received tumor implants in the 4^th^ mammary gland of right flank. Randomization to experimental groups with comparable mean tumor volumes was performed, when the tumors reached over 70 mm^3^ volume on average. The day of randomization was designated as day 0 of the experiment and the dosing was started on day 0. Tumor volumes and relative tumor volumes were calculated as with the MAXFTN-MX1 model. Tumor volumes were determined three times a week and body weight at least once per week. The ethical limit for the tumor size was 2 cm. According to the ethical endpoints, the mice were terminated if their body weight loss was ≥15% or if their body condition scoring decreased to <2.

The mice in MAXFTN-MX1 and HCI-002 experiment were treated as follows. Group 1 received vehicle 1 (5 % DMSO, 95 % Natrosol-1 % solution) once a day and vehicle 2 (20 % Solutol HS15 / 20 % SB-b-CD (Captisol) / 50 mM phosphate buffer, pH 7.4) twice a day. Group 2 was treated with 10mg/kg mubritinib in vehicle 1 once a day and received also vehicle 2 twice a day. Group 3 was treated with 200mg/kg CB-839 in vehicle 2 twice a day and received also vehicle 1 once a day. Group 4 was treated with mubritinib once a day and CB-839 twice a day. The treatments were administered by i.g. route for 21 days in the MAXFTN-MX1. In the HCI-002 experiment, the treatments were administered via i.g. route for 11. The drug administrations were discontinued after 11 days of treatment due to vehicle 2-caused diarrhea in the NSG mice. The *in vivo* experiments were not blinded; bias could only be introduced by TV measurement, but this can be considered negligible in comparison to the overall biological variation of tumor growth.

### Statistical analysis

Statistical analyses were performed using GraphPad Prism 9 and R. Data are presented as the mean ± s.d. Two-tailed t-test was used for comparing two groups. Two-way ANOVA followed by Šidák’s or Dunnett’s comparison tests were used to compare multiple variables. One-tailed Mann-Whitney test used to compare relative PDX tumor volumes in MAXFTN-MX1 model and unpaired t-test with Welch’s correction was used compare relative PDX tumor volumes in HCI-002 model. Log-rank (Mantel-Cox) test was used to test the significance of survival of PDXs. Statistical significance was defined as *p < 0.05, **p < 0.01, ***p < 0.001 and ****p < 0.0001.

## Data availability

All data associated with this study are present in the paper or the supplementary materials. RNAseq gene expression counts can be found in Supplementary Data 1. All metabolites and their isotopogues for figure 5B can be found in Supplementary Data 2.

## ACKNOWLEDGEMENTS

We thank HiLIFE and Biocenter Finland funded Institute for Molecular Medicine Finland Metabolomics Unit (FIMM Meta), Biomedicum Imaging Unit (BIU), Biomedicum Functional Genomics Unit (FuGU) and Finnish Genome editing center (FINGEEC) for their services. Drug sensitivity screening was carried out at the FIMM High Throughput Biomedicine Unit, which is hosted by the University of Helsinki and supported by HiLIFE and Biocenter Finland. We thank Tiina Raatikainen, Katriina Karjalainen, Elina Hurskainen and Ellisiv Nyhamar for their technical support. We thank Tiina Raatikainen also for the help with the HCI-002 mouse experiment. We acknowledge Dr. Alana Welm at Huntsman Cancer Institute, University of Utah for HCI-002 PDX tumor material. We thank Charles River Laboratories, Germany for executing the patient-derived xenograft mouse experiment with MAXFTN-MX1 model. Schematic images were created with BioRender.com. This work was funded by the Academy of Finland, Business Finland, the Finnish Cancer Organizations, the Sigrid Juselius Foundation, Jane and Aatos Erkko Foundation, Breast Cancer Now (2017NovPCC1067) and RESCUER project, which has received funding from the European Union’s Horizon 2020 research and innovation programme under grant agreement No. 847912. This work was also supported by the CDMRP W81XWH211-0773/-0774. Opinions, interpretations, conclusions, and recommendations are those of the author and are not necessarily endorsed by the Department of Defense. J.M.A. was funded by Integrative Life Sciences Doctoral Program, The University of Helsinki Doctoral School, Biomedicum Helsinki Foundation, Emil Aaltonen Foundation, Ida Montin Foundation, The Maud Kuistila Memorial Foundation and Cancer Foundation Finland. M.S was funded by Doctoral Programme in Biomedicine, The University of Helsinki Doctoral School, Orion Research Foundation, The Paulo Foundation and Cancer Foundation Finland. R.A. was funded by a Doctoral fellowship of the Integrated Life Sciences Graduate School. T.A. was supported by Academy of Finland (grants 340141, 344698, and 345803), the Cancer Society of Finland, the Norwegian Cancer Society, the Sigrid Jusélius Foundation, and iCAN – Digital Precision Cancer Medicine Flagship (iCAN-MULTIDRUG). C.B.J. was supported by Academy of Finland (336455) and Jane and Aatos Erkko Foundation.

## AUTHOR CONTRIBUTIONS

J.M.A., M.S. and J.K. designed the concept of the study. J.M.A., M.S., P.M.M., L.I., A.O.H., J.A., R.A. B.P., M.P., H.A.H., P.G., M.V., T.A.T., K.S., R.D. and M.V.R.P. performed the experiments. J.M.A. and M.S. completed the statistical analysis for biological experiments. J.S., D.N. and A.P. performed bioinformatic analysis. J.M.A., J.K. and M.S. wrote the manuscript. K.W., M.A.H., A.G., J.W., T.A., C.B.J., A.I.N. and J.K. supervised the studies. M.M., P.K., L.N., T.M., J.M. and P.H. contributed to clinical data/analysis.

## REFERENCES

1. Blackwood, E. M. & Eisenman, R. N. Max: A Helix-Loop-Helix Zipper Protein That Forms a Sequence-Specific DNA-Binding Complex with Myc. Science (1979) 251, 1211–1217 (1991).

2. Dang, C. V. MYC on the Path to Cancer. Cell 149, 22–35 (2012).

3. Horiuchi, D. et al. MYC pathway activation in triple-negative breast cancer is synthetic lethal with CDK inhibition. Journal of Experimental Medicine 209, 679–696 (2012).

4. Naidu, R., Wahab Abdul, N., Yadav, M. & Kutty Kannan, M. Protein expression and molecular analysis of c-myc gene in primary breast carcinomas using immunohistochemistry and differential polymerase chain reaction. Int J Mol Med 9, 189–196 (2002).

5. Haikala, H. M. et al. Pharmacological reactivation of MYC-dependent apoptosis induces susceptibility to anti-PD-1 immunotherapy. Nat Commun 10, (2019).

6. Koboldt, D. C. et al. Comprehensive molecular portraits of human breast tumours. Nature 490, 61–70 (2012).

7. Green, A. R. et al. MYC functions are specific in biological subtypes of breast cancer and confers resistance to endocrine therapy in luminal tumours. Br J Cancer 114, 917– 928 (2016).

8. Donati, G. & Amati, B. MYC and therapy resistance in cancer: risks and opportunities. Mol Oncol 16, 3828–3854 (2022).

9. Morrish, F., Neretti, N., Sedivy, J. M. & Hockenbery, D. M. The oncogene c-Myc coordinates regulation of metabolic networks to enable rapid cell cycle entry. Cell Cycle 7, 1054–1066 (2008).

10. Evan, G. I. et al. Induction of apoptosis in fibroblasts by c-myc protein. Cell 69, 119– 128 (1992).

11. Nieminen, A. I., Partanen, J. I., Hau, A. & Klefstrom, J. c-Myc primed mitochondria determine cellular sensitivity to TRAIL-induced apoptosis. EMBO J 26, 1055–1067 (2007).

12. Nieminen, A. I. et al. Myc-induced AMPK-phospho p53 pathway activates Bak to sensitize mitochondrial apoptosis. Proc Natl Acad Sci U S A 110, E1839–48 (2013).

13. Lee, J. V et al. Combinatorial immunotherapies overcome MYC-driven immune evasion in triple negative breast cancer. Nat Commun 13, 3671 (2022).

14. Pourdehnad, M. et al. Myc and mTOR converge on a common node in protein synthesis control that confers synthetic lethality in Myc-driven cancers. Proceedings of the National Academy of Sciences 110, 11988–11993 (2013).

15. Yang, D. et al. Therapeutic potential of a synthetic lethal interaction between the MYC proto-oncogene and inhibition of aurora-B kinase. Proceedings of the National Academy of Sciences 107, 13836–13841 (2010).

16. Xiao, D. et al. Polo-like Kinase-1 Regulates Myc Stabilization and Activates a Feedforward Circuit Promoting Tumor Cell Survival. Mol Cell 64, 493–506 (2016).

17. Wise, D. R. et al. Myc regulates a transcriptional program that stimulates mitochondrial glutaminolysis and leads to glutamine addiction. Proc Natl Acad Sci U S A 105, 18782–18787 (2008).

18. Alborzinia, H. et al. MYCN mediates cysteine addiction and sensitizes neuroblastoma to ferroptosis. Nat Cancer 3, 471–485 (2022).

19. Muir, A. et al. Environmental cystine drives glutamine anaplerosis and sensitizes cancer cells to glutaminase inhibition. Elife 6, e27713 (2017).

20. Le, A. et al. Glucose-independent glutamine metabolism via TCA cycling for proliferation and survival in B cells. Cell Metab 15, 110–121 (2012).

21. Yoo, H. C., Yu, Y. C., Sung, Y. & Han, J. M. Glutamine reliance in cell metabolism. Exp Mol Med 52, 1496–1516 (2020).

22. DeBerardinis, R. J. et al. Beyond aerobic glycolysis: Transformed cells can engage in glutamine metabolism that exceeds the requirement for protein and nucleotide synthesis. Proceedings of the National Academy of Sciences 104, 19345–19350 (2007).

23. Chien, A. J. et al. A phase Ib trial of the cyclin-dependent kinase inhibitor dinaciclib (dina) in combination with pembrolizumab (P) in patients with advanced triple-negative breast cancer (TNBC) and response correlation with MYC-overexpression. Journal of Clinical Oncology 38, 1076 (2020).

24. Jung, M. et al. A Myc Activity Signature Predicts Poor Clinical Outcomes in Myc-Associated Cancers. Cancer Res 77, 971–981 (2017).

25. Chandriani, S. et al. A Core MYC Gene Expression Signature Is Prominent in Basal-Like Breast Cancer but Only Partially Overlaps the Core Serum Response. PLoS One 4, e6693 (2009).

26. Kreuzaler, P. et al. Vitamin B5 supports MYC oncogenic metabolism and tumor progression in breast cancer. Nat Metab 5, 1870–1886 (2023).

27. Dent, R. et al. Triple-Negative Breast Cancer: Clinical Features and Patterns of Recurrence. Clinical Cancer Research 13, 4429–4434 (2007).

28. Surveillance Research Program, N. C. I. SEER*Explorer: An interactive website for SEER cancer statistics. https://seer.cancer.gov/statistics-network/explorer/ (2023).

29. Nusinow, D. P. et al. Quantitative Proteomics of the Cancer Cell Line Encyclopedia. Cell 180, 387–402.e16 (2020).

30. Mertins, P. et al. Proteogenomics connects somatic mutations to signalling in breast cancer. Nature 534, 55–62 (2016).

31. Liberzon, A. et al. The Molecular Signatures Database Hallmark Gene Set Collection. Cell Syst 1, 417–425 (2015).

32. Morrish, F. & Hockenbery, D. MYC and Mitochondrial Biogenesis. Cold Spring Harbor Perspectives in Medicine 4, (2014).

33. Rath, S. et al. MitoCarta3.0: an updated mitochondrial proteome now with sub-organelle localization and pathway annotations. Nucleic Acids Res 49, D1541–D1547 (2021).

34. Liberzon, A. et al. Molecular signatures database (MSigDB) 3.0. Bioinformatics 27, 1739–1740 (2011).

35. Jassal, B. et al. The reactome pathway knowledgebase. Nucleic Acids Res 48, D498– D503 (2020).

36. Li, F. et al. Myc Stimulates Nuclearly Encoded Mitochondrial Genes and Mitochondrial Biogenesis. Mol Cell Biol 25, 6225–6234 (2005).

37. Bridges, H. R., Jones, A. J. Y., Pollak, M. N. & Hirst, J. Effects of metformin and other biguanides on oxidative phosphorylation in mitochondria. Biochemical Journal 462, 475–487 (2014).

38. Wheaton, W. W. et al. Metformin inhibits mitochondrial complex I of cancer cells to reduce tumorigenesis. Elife 3, e02242 (2014).

39. Narita, M. et al. Rb-Mediated Heterochromatin Formation and Silencing of E2F Target Genes during Cellular Senescence. Cell 113, 703–716 (2003).

40. Lee, S. & Schmitt, C. A. The dynamic nature of senescence in cancer. Nature Cell Biology 2019 21:1 21, 94–101 (2019).

41. Ashraf, H. M., Fernandez, B. & Spencer, S. L. The intensities of canonical senescence biomarkers integrate the duration of cell-cycle withdrawal. Nature Communications 2023 14:1 14, 1–13 (2023).

42. Baccelli, I. et al. Mubritinib Targets the Electron Transport Chain Complex I and Reveals the Landscape of OXPHOS Dependency in Acute Myeloid Leukemia. Cancer Cell 36, 84–99.e8 (2019).

43. Ellinghaus, P. et al. BAY 87-2243, a highly potent and selective inhibitor of hypoxia-induced gene activation has antitumor activities by inhibition of mitochondrial complex I. Cancer Med 2, 611–624 (2013).

44. Chung, I. et al. Cork-in-bottle mechanism of inhibitor binding to mammalian complex I. Sci Adv 7, eabg4000 (2021).

45. Osthus, R. C. et al. Deregulation of glucose transporter 1 and glycolytic gene expression by c-Myc. Journal of Biological Chemistry 275, 21797–21800 (2000).

46. Kim, J. et al. Evaluation of Myc E-Box Phylogenetic Footprints in Glycolytic Genes by Chromatin Immunoprecipitation Assays. Mol Cell Biol 24, 5923–5936 (2004).

47. Oliynyk, G. et al. MYCN-enhanced Oxidative and Glycolytic Metabolism Reveals Vulnerabilities for Targeting Neuroblastoma. iScience 21, 188–204 (2019).

48. Lee, K. min et al. MYC and MCL1 Cooperatively Promote Chemotherapy-Resistant Breast Cancer Stem Cells via Regulation of Mitochondrial Oxidative Phosphorylation. Cell Metab 26, 633–647.e7 (2017).

49. Donati, G. et al. Targeting mitochondrial respiration and the BCL2 family in high-grade MYC-associated B-cell lymphoma. Mol Oncol 16, 1132–1152 (2022).

50. Caro, P. et al. Metabolic Signatures Uncover Distinct Targets in Molecular Subsets of Diffuse Large B Cell Lymphoma. Cancer Cell 22, 547–560 (2012).

51. Vazquez, F. et al. PGC1α Expression Defines a Subset of Human Melanoma Tumors with Increased Mitochondrial Capacity and Resistance to Oxidative Stress. Cancer Cell 23, 287–301 (2013).

52. Shang, Y. et al. Overexpression of UQCRC2 is correlated with tumor progression and poor prognosis in colorectal cancer. Pathol Res Pract 214, 1613–1620 (2018).

53. Janiszewska, M. et al. Imp2 controls oxidative phosphorylation and is crucial for preserving glioblastoma cancer stem cells. Genes Dev 26, 1926–1944 (2012).

54. Yang, Y. & Sauve, A. A. NAD(+) metabolism: Bioenergetics, signaling and manipulation for therapy. Biochim Biophys Acta 1864, 1787–1800 (2016).

55. Stein, L. R. & Imai, S. The dynamic regulation of NAD metabolism in mitochondria. Trends in Endocrinology & Metabolism 23, 420–428 (2012).

56. DeBerardinis, R. J. & Chandel, N. S. Fundamentals of cancer metabolism. Sci Adv 2, e1600200 (2016).

57. Gan, L. et al. Metabolic targeting of oncogene MYC by selective activation of the proton-coupled monocarboxylate family of transporters. Oncogene 35, 3037–3048 (2016).

58. Yuneva, M., Zamboni, N., Oefner, P., Sachidanandam, R. & Lazebnik, Y. Deficiency in glutamine but not glucose induces MYC-dependent apoptosis in human cells. J Cell Biol 178, 93–105 (2007).

59. Anderton, B. et al. MYC-driven inhibition of the glutamate-cysteine ligase promotes glutathione depletion in liver cancer. EMBO Rep 18, 569–585 (2017).

60. Donati, G. et al. Oxidative stress enhances the therapeutic action of a respiratory inhibitor in MYC -driven lymphoma . EMBO Mol Med 15, (2023).

61. Zhang, X. & Dang, C. V. Time to hit pause on mitochondria-targeting cancer therapies. Nat Med 29, 29–30 (2023).

62. Takeda. Safety and Tolerability Study of TAK-165 in Subjects With Tumors Expressing HER2, NCT00034281. https://classic.clinicaltrials.gov/ct2/show/NCT00034281 (2012).

63. Dowlati, A. et al. Novel Phase I Dose De-escalation Design Trial to Determine the Biological Modulatory Dose of the Antiangiogenic Agent SU5416. Clinical Cancer Research 11, 7938–7944 (2005).

64. Barretina, J. et al. The Cancer Cell Line Encyclopedia enables predictive modelling of anticancer drug sensitivity. Nature 483, 603–607 (2012).

65. Robinson, M. D., McCarthy, D. J. & Smyth, G. K. edgeR: a Bioconductor package for differential expression analysis of digital gene expression data. Bioinformatics 26, 139–140 (2010).

66. Subramanian, A. et al. Gene set enrichment analysis: A knowledge-based approach for interpreting genome-wide expression profiles. Proceedings of the National Academy of Sciences 102, 15545 LP – 15550 (2005).

67. Wu, D. & Smyth, G. K. Camera: a competitive gene set test accounting for inter-gene correlation. Nucleic Acids Res 40, e133–e133 (2012).

68. Pal, B. et al. A single-cell RNA expression atlas of normal, preneoplastic and tumorigenic states in the human breast. EMBO J 40, (2021).

69. Wu, S. Z. et al. A single-cell and spatially resolved atlas of human breast cancers. Nature Genetics 2021 53:9 53, 1334–1347 (2021).

70. Bassiouni, R. et al. Spatial Transcriptomic Analysis of a Diverse Patient Cohort Reveals a Conserved Architecture in Triple-Negative Breast Cancer. Cancer Res 83, 34–48 (2023).

71. Hao, Y. et al. Dictionary learning for integrative, multimodal and scalable single-cell analysis. Nature Biotechnology 2023 42:2 42, 293–304 (2023).

72. Hao, Y. et al. Integrated analysis of multimodal single-cell data. Cell 184, 3573–3587.e29 (2021).

73. Stuart, T. et al. Comprehensive Integration of Single-Cell Data. Cell 177, 1888–1902.e21 (2019).

74. Butler, A., Hoffman, P., Smibert, P., Papalexi, E. & Satija, R. Integrating single-cell transcriptomic data across different conditions, technologies, and species. Nature Biotechnology 2018 36:5 36, 411–420 (2018).

75. Satija, R., Farrell, J. A., Gennert, D., Schier, A. F. & Regev, A. Spatial reconstruction of single-cell gene expression data. Nature Biotechnology 2015 33:5 33, 495–502 (2015).

76. Hafemeister, C. & Satija, R. Normalization and variance stabilization of single-cell RNA-seq data using regularized negative binomial regression. Genome Biol 20, 1–15 (2019).

77. Choudhary, S. & Satija, R. Comparison and evaluation of statistical error models for scRNA-seq. Genome Biol 23, 1–20 (2022).

78. Zappia, L. & Oshlack, A. Clustering trees: a visualization for evaluating clusterings at multiple resolutions. Gigascience 7, 1–9 (2018).

79. Yoshihara, K. et al. Inferring tumour purity and stromal and immune cell admixture from expression data. Nature Communications 2013 4:1 4, 1–11 (2013).

80. Awadhpersad, R. & Jackson, C. B. High-Resolution Respirometry to Assess Bioenergetics in Cells and Tissues Using Chamber- and Plate-Based Respirometers. JoVE e63000 (2021) doi:doi:10.3791/63000.

81. Jackson, C. B., Gallati, S. & Schaller, A. qPCR-based mitochondrial DNA quantification: Influence of template DNA fragmentation on accuracy. Biochem Biophys Res Commun 423, 441–447 (2012).

82. Rooney, J. P. et al. PCR Based Determination of Mitochondrial DNA Copy Number in Multiple Species. Methods in Molecular Biology 1241, 23–38 (2015).

83. Vandesompele, J. et al. Accurate normalization of real-time quantitative RT-PCR data by geometric averaging of multiple internal control genes. Genome Biol 3, 1–12 (2002).

84. Stirling, D. R. et al. CellProfiler 4: improvements in speed, utility and usability. BMC Bioinformatics 22, 433 (2021).

85. Wilks, C. et al. recount3: summaries and queries for large-scale RNA-seq expression and splicing. Genome Biol 22, 323 (2021).

86. Lehmann, B. D. et al. Identification of human triple-negative breast cancer subtypes and preclinical models for selection of targeted therapies. J Clin Invest 121, 2750–2767 (2011).

87. Gautam, P. et al. Identification of selective cytotoxic and synthetic lethal drug responses in triple negative breast cancer cells. Mol Cancer 15, 34 (2016).

88. Kulesskiy, E., Saarela, J., Turunen, L. & Wennerberg, K. Precision Cancer Medicine in the Acoustic Dispensing Era: Ex Vivo Primary Cell Drug Sensitivity Testing. J Lab Autom 21, 27–36 (2016).

89. Chen, Y. et al. Robust scoring of selective drug responses for patient-tailored therapy selection. Nat Protoc 19, 60–82 (2024).

90. Potdar, S. et al. Breeze 2.0: an interactive web-tool for visual analysis and comparison of drug response data. Nucleic Acids Res 51, W57–W61 (2023).

