## Supplementary information for "Complex I Drives Glutamine-Dependent TCA Cycle to Support Viability of MYC^high^ Breast Cancer Cells"

**Supplementary Figure 1.**

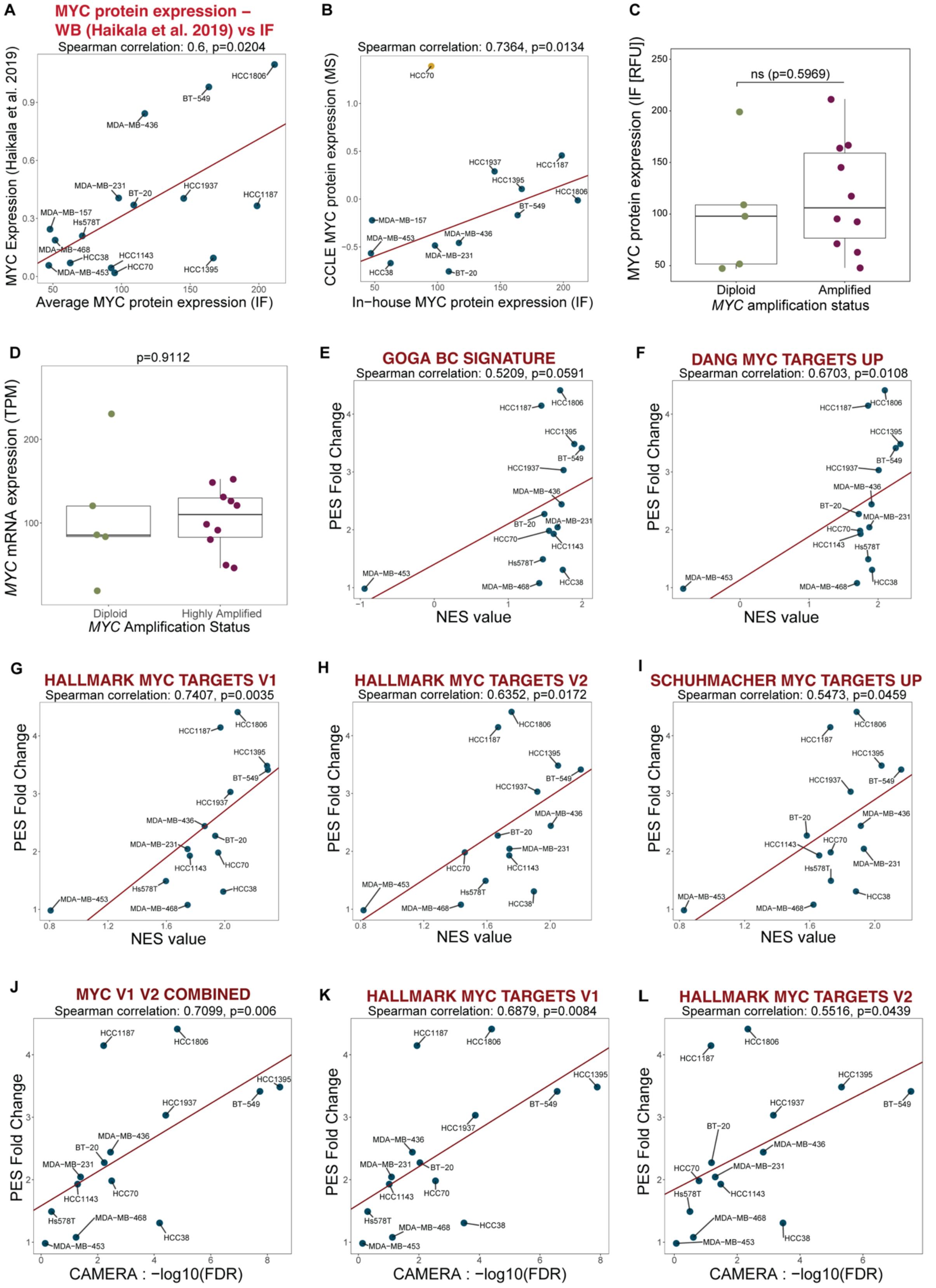


**Supplementary Figure 1. Correlation between MYC protein expression and MYC activity signatures. A** Correlation between MYC protein expression levels estimated from the high content immunofluorescence (Fig. 1A) and western blot analyses (in [5]). **B** Correlation between MYC protein expression levels estimated from the high content immunofluorescence and CCLE mass spectrometry analyses. HCC70 cell line was left out from the correlation analysis as an outlier (yellow dot). **C** Association between *MYC* amplification (obtained from CCLE) and the MYC protein expression levels in TNBC cells analyzed using linear regression **D** Statistical association analysis (linear regression) between the *MYC* mRNA expression (Transcripts Per Kilobase Million/TMP) and *MYC* amplification status (obtained from CCLE) in TNBC cells. **E-I** Spearman correlation between the PES and the five clustered MYC activity gene sets (from Fig. 1D) in TNBC cells. **J-L** Spearman correlation between the PES and MYC V1V2 (**I**), HALLMARK_MYC_TARGETS_V1 (**J**) or HALLMARK_MYC_TARGETS_V2 (**K**) in TNBC cells.

**Supplementary Figure 2.**

**A**

CCLE RNAseq data

| Gene set name | Pearson_r | FDR |
|---|---|---|
| HP_ABNORMAL_ACTIVITY_OF_MITOCHONDRIAL_RESPIRATORY_CHAIN | 0.66 | <0.0001 |
| OXPHOS Complex I assembly factors | 0.64 | <0.0001 |
| OXPHOS assembly factors | 0.63 | <0.0001 |
| GOBP_MITOCHONDRIAL_RESPIRATORY_CHAIN_COMPLEX_ASSEMBLY | 0.59 | <0.0001 |
| OXPHOS Complex IV assembly factors | 0.54 | <0.0001 |
| OXPHOS | 0.52 | <0.0001 |
| OXPHOS Complex I | 0.51 | <0.0001 |
| WP_MITOCHONDRIAL_COMPLEX_I_ASSEMBLY_MODEL_OXPHOS_SYSTEM | 0.50 | <0.0001 |
| REACTOME_RESPIRATORY_ELECTRON_TRANSPORT_ATP_SYNTHESIS_BY _CHEMIOSMOTIC_COUPLING_AND_HEAT_PRODUCTION_BY_UNCOUPLING _PROTEINS | 0.49 | <0.0001 |
| REACTOME_RESPIRATORY_ELECTRON_TRANSPORT | 0.48 | <0.0001 |
| OXPHOS Complex IV | 0.42 | <0.0001 |
| REACTOME_THE_CITRIC_ACID_TCA_CYCLE_AND_RESPIRATORY_ELECTR ON_TRANSPORT | 0.41 | <0.0001 |
| HALLMARK_OXIDATIVE_PHOSPHORYLATION | 0.39 | <0.0001 |
| WP_ELECTRON_TRANSPORT_CHAIN_OXPHOS_SYSTEM_IN_MITOCHONDRIA | 0.34 | <0.0001 |
| OXPHOS Complex I subunits | 0.33 | <0.0001 |
| GOBP_OXIDATIVE_PHOSPHORYLATION | 0.32 | <0.0001 |
| GOBP_ATP_SYNTHESIS_COUPLED_ELECTRON_TRANSPORT | 0.31 | <0.0001 |
| GOBP_RESPIRATORY_ELECTRON_TRANSPORT_CHAIN | 0.31 | <0.0001 |
| OXPHOS subunits | 0.28 | <0.0001 |
| GOBP_MITOCHONDRIAL_ELECTRON_TRANSPORT_NADH_TO_UBIQUINONE | 0.22 | <0.0001 |
| WP_OXIDATIVE_PHOSPHORYLATION | 0.21 | <0.0001 |
| GOBP_REGULATION_OF_OXIDATIVE_PHOSPHORYLATION | -0.19 | <0.0001 |
| GOBP_ELECTRON_TRANSPORT_CHAIN | 0.13 | <0.0001 |
| KEGG_OXIDATIVE_PHOSPHORYLATION | 0.11 | 0.00046 |
| GOBP_MITOCHONDRIAL_ELECTRON_TRANSPORT_CYTOCHROME_C_TO_ OXYGEN | 0.07 | 0.02229 |
| OXPHOS Complex IV subunits | -0.07 | 0.03395 |

**B**

TCGA TNBC RNAseq data

| Gene set name | Pearson's r | FDR |
|---|---|---|
| OXPHOS Complex IV assembly factors | 0.57 | <0.0001 |
| OXPHOS Complex I assembly factors | 0.55 | <0.0001 |
| OXPHOS assembly factors | 0.54 | <0.0001 |
| HP_ABNORMAL_ACTIVITY_OF_MITOCHONDRIAL_RESPIRATORY_CHAIN | 0.53 | <0.0001 |
| GOBP_MITOCHONDRIAL_RESPIRATORY_CHAIN_COMPLEX_ASSEMBLY | 0.43 | <0.0001 |
| OXPHOS Complex I | 0.43 | <0.0001 |
| OXPHOS | 0.42 | <0.0001 |
| WP_MITOCHONDRIAL_COMPLEX_I_ASSEMBLY_MODEL_OXPHOS_SYST EM | 0.41 | <0.0001 |
| REACTOME_THE_CITRIC_ACID_TCA_CYCLE_AND_RESPIRATORY_ELEC TRON_TRANSPORT | 0.39 | <0.0001 |
| OXPHOS Complex IV | 0.39 | 0.00013 |
| REACTOME_RESPIRATORY_ELECTRON_TRANSPORT | 0.38 | 0.00013 |
| REACTOME_RESPIRATORY_ELECTRON_TRANSPORT_ATP_SYNTHESIS _BY_CHEMIOSMOTIC_COUPLING_AND_HEAT_PRODUCTION_BY_UNCO UPLING_PROTEINS | 0.38 | 0.00015 |
| HALLMARK_OXIDATIVE_PHOSPHORYLATION | 0.34 | 0.00075 |
| GOBP_OXIDATIVE_PHOSPHORYLATION | 0.33 | 0.00127 |
| GOBP_ATP_SYNTHESIS_COUPLED_ELECTRON_TRANSPORT | 0.32 | 0.00127 |
| OXPHOS Complex I subunits | 0.31 | 0.00237 |
| GOBP_RESPIRATORY_ELECTRON_TRANSPORT_CHAIN | 0.30 | 0.00272 |
| WP_ELECTRON_TRANSPORT_CHAIN_OXPHOS_SYSTEM_IN_MITOCHON DRIA | 0.28 | 0.00661 |
| WP_OXIDATIVE_PHOSPHORYLATION | 0.26 | 0.00977 |
| OXPHOS subunits | 0.24 | 0.02020 |
| KEGG_OXIDATIVE_PHOSPHORYLATION | 0.21 | 0.03621 |
| GOBP_REGULATION_OF_OXIDATIVE_PHOSPHORYLATION | -0.19 | 0.05944 |
| GOBP_MITOCHONDRIAL_ELECTRON_TRANSPORT_NADH_TO_UBIQUIN ONE | 0.18 | 0.07160 |
| OXPHOS Complex IV subunits | -0.14 | 0.17357 |
| GOBP_ELECTRON_TRANSPORT_CHAIN | 0.10 | 0.31269 |
| GOBP_MITOCHONDRIAL_ELECTRON_TRANSPORT_CYTOCHROME_C_T O_OXYGEN | 0.08 | 0.43805 |

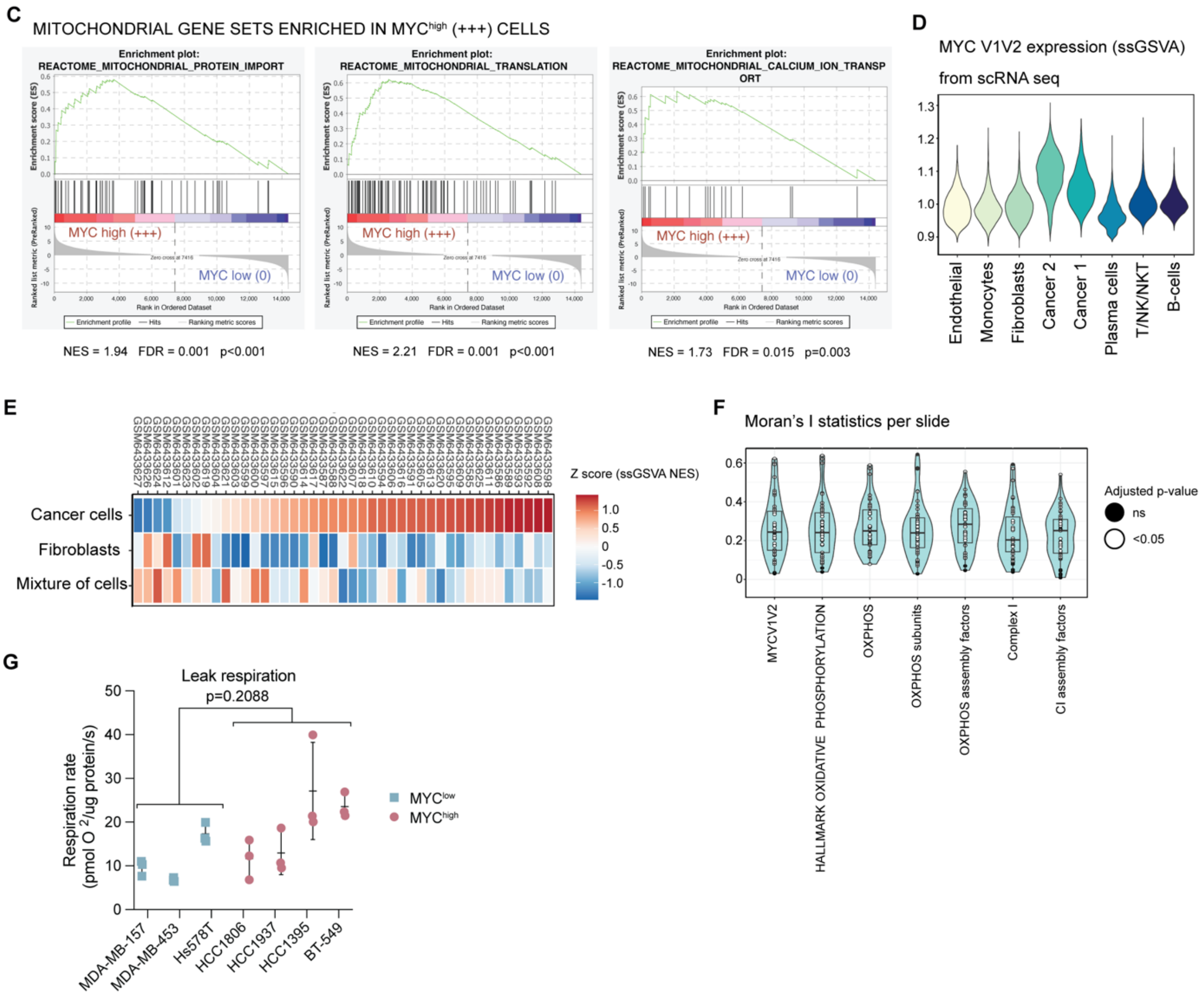


**Supplementary Figure 2. Mitochondrial respiration processes in MYC$^{high}$ and MYC$^{low}$ TNBC. A** Pearson correlation coefficient values, r, and corresponding FDR values for Pearson correlations between enrichment scores (ES) of mitochondrial respiration gene sets and MYC

V1V2 ES. ESs were calculated from CCLE RNAseq data. **B** Pearson correlation coefficient values, r, and corresponding FDR values for Pearson correlations between ES of mitochondrial respiration gene sets and MYC V1V2 ES. ESs were calculated from TCGA TNBC sample RNAseq data. **C** Mitochondrial gene sets enriched in MYC$^{high}$ (+++) *versus* MYC$^{low}$ (0). **D** MYC V1V2 enrichment scores from scRNA sequencing in different cell types. **E** MYC V1V2 enrichment score calculated per spot and cell type for each slide with ssGSVA. All results were scaled slide-wise. **F** Moran's I statistics calculated per patient. **G** Leak respiration measured by Oroboros.

**Supplementary Figure 3.**

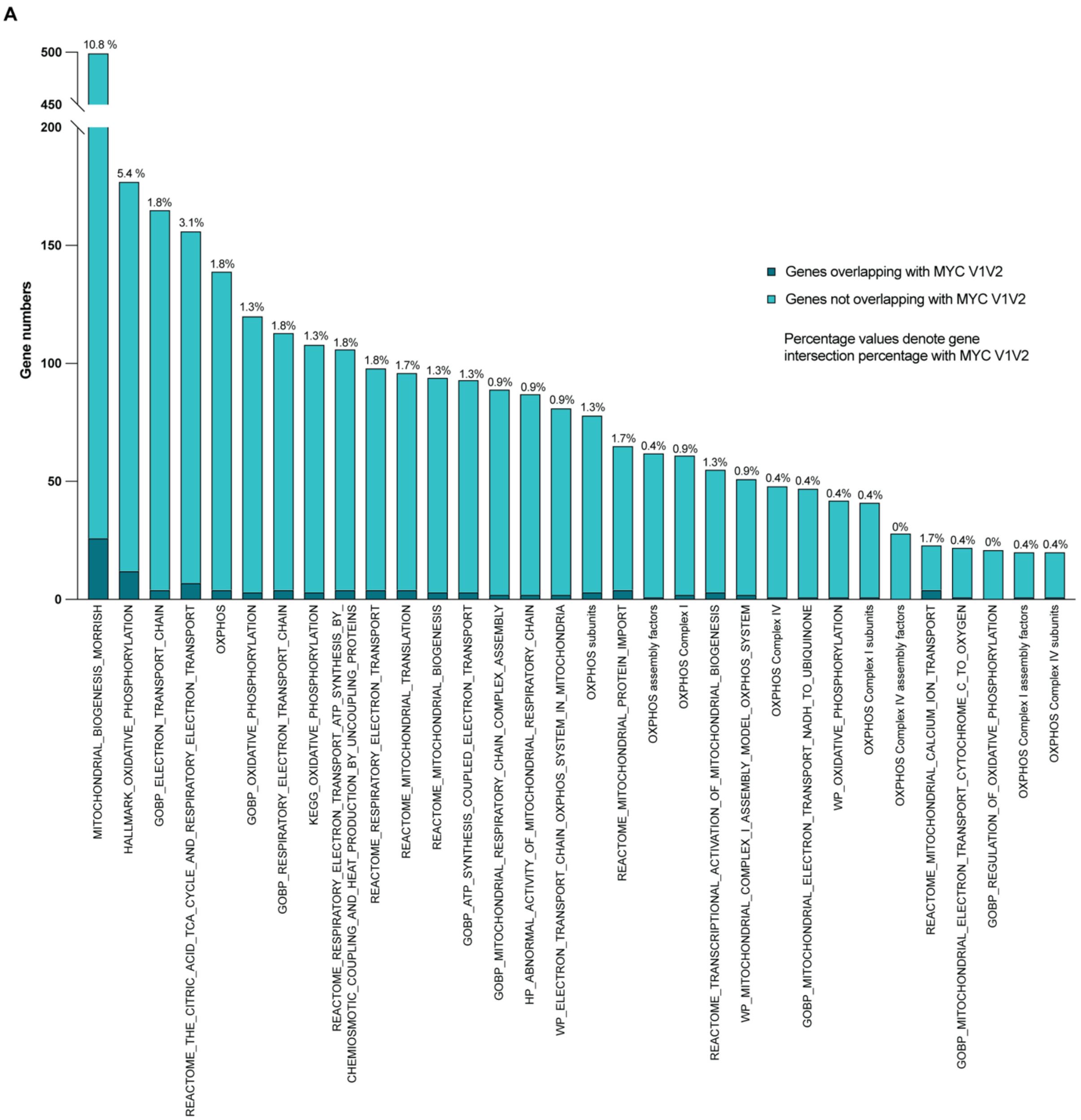


**Supplementary Figure 3. Overlap between mitochondrial gene sets and MYC V1V2 gene set. A** The overlap between the mitochondrial gene sets used in the study and MYC V1V2 gene set shown both in numbers of the genes and percentages denoting the gene intersection percentage between each gene set and MYC V1V2 gene set.

**Supplementary Figure 4.**

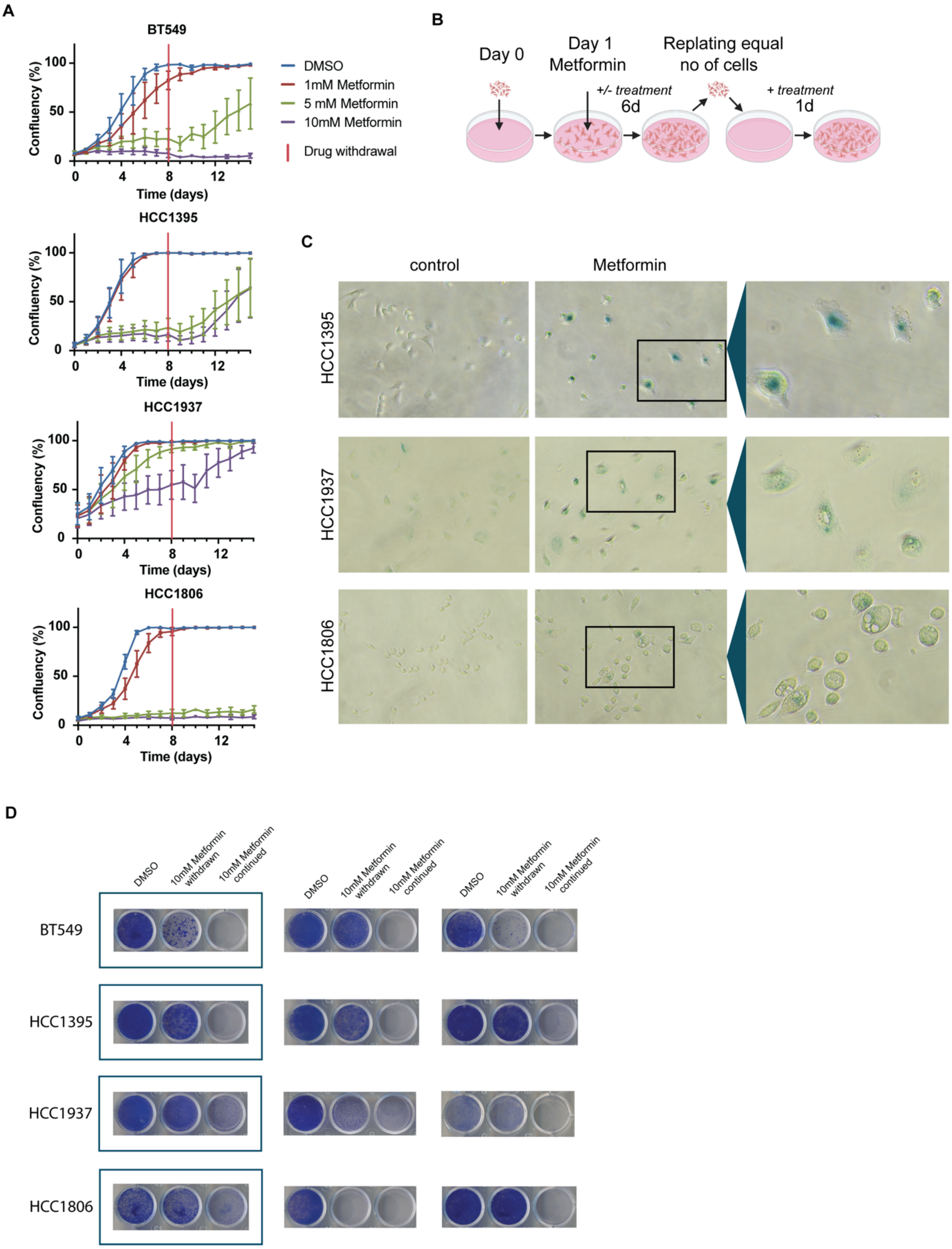


**Supplementary Figure 4. Metformin withdrawal and replating experiments. A** Metformin withdrawal on MYC$^{high}$ TNBC cells in time-lapse video microscopy experiment. The cells were treated with metformin 8 days before the drug withdrawal and in total imaged for 15 days with

IncuCyte. **B** Schematic illustration of senescence-associated beta-galactosidase activity replating experiment. **C** Senescence-associated beta-galactosidase activity in metformin-treated MYC$^{high}$ TNBC cells. **D** Repeats of replating assay for senescense (Fig. 3F).

**Supplementary Figure 5.**

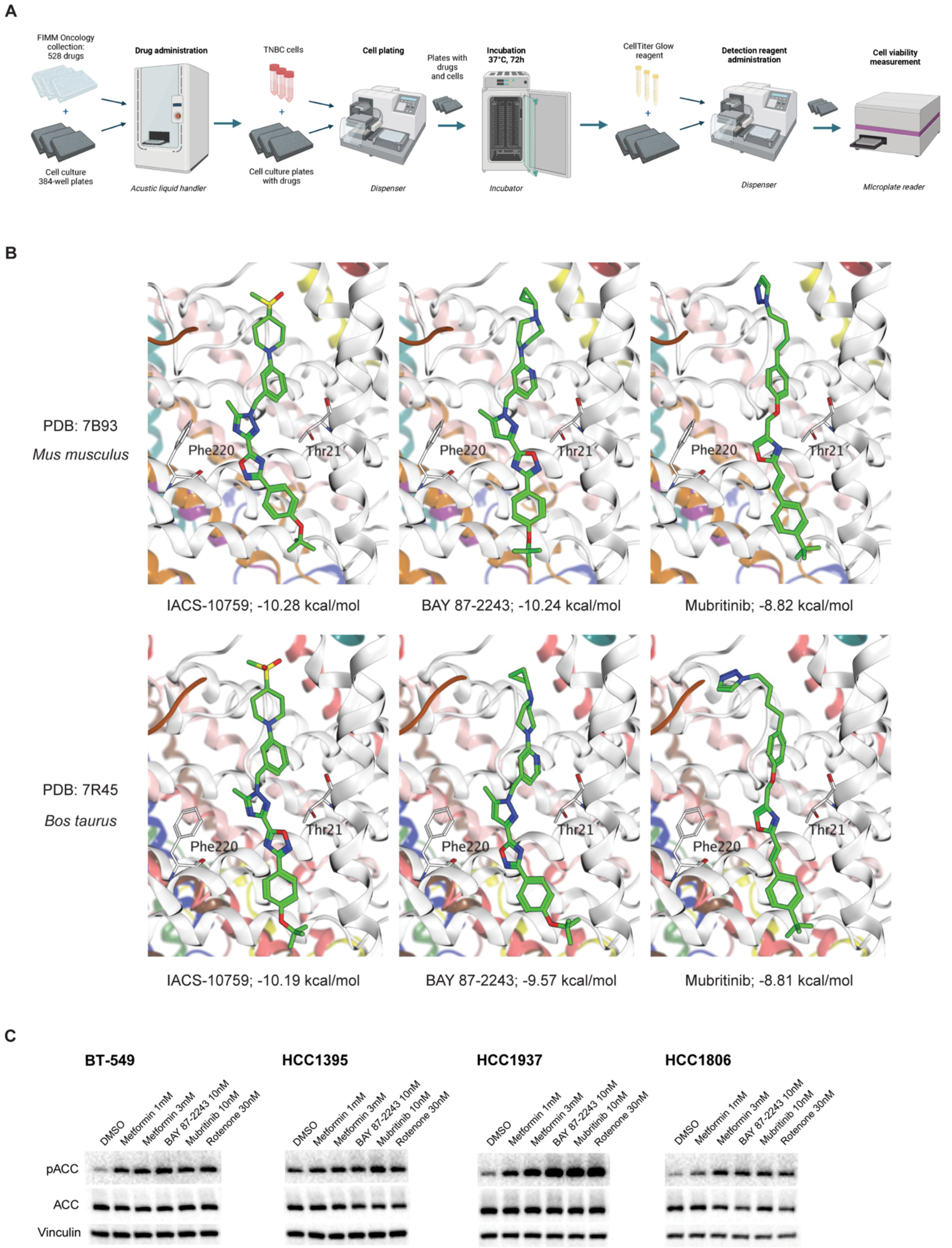


**Supplementary Figure 5. Drug sensitivity testing. A** Detailed workflow for the drug sensitivity testing. **B** Comparative molecular docking analysis of IACS-10759, BAY 87-2243 and Mubritinib binding to Q-site of complex I. Two cryo-EM structures of mitochondrial complex I bound to IACS-2858 (PDB ID:7B93) and phenformin-derivative IM1761092 (PDB ID:7R45) were used as protein templates. The binding affinity (kcal/mol), calculated by the scoring function, is presented as a computational estimate of the stability of the protein-ligand complex. **C** Western blot analysis for pACC and ACC in $MYC^{high}$ cells after complex I inhibitor treatment. Representative stainings shown from three biological replicates.

**Supplementary Figure 6.**

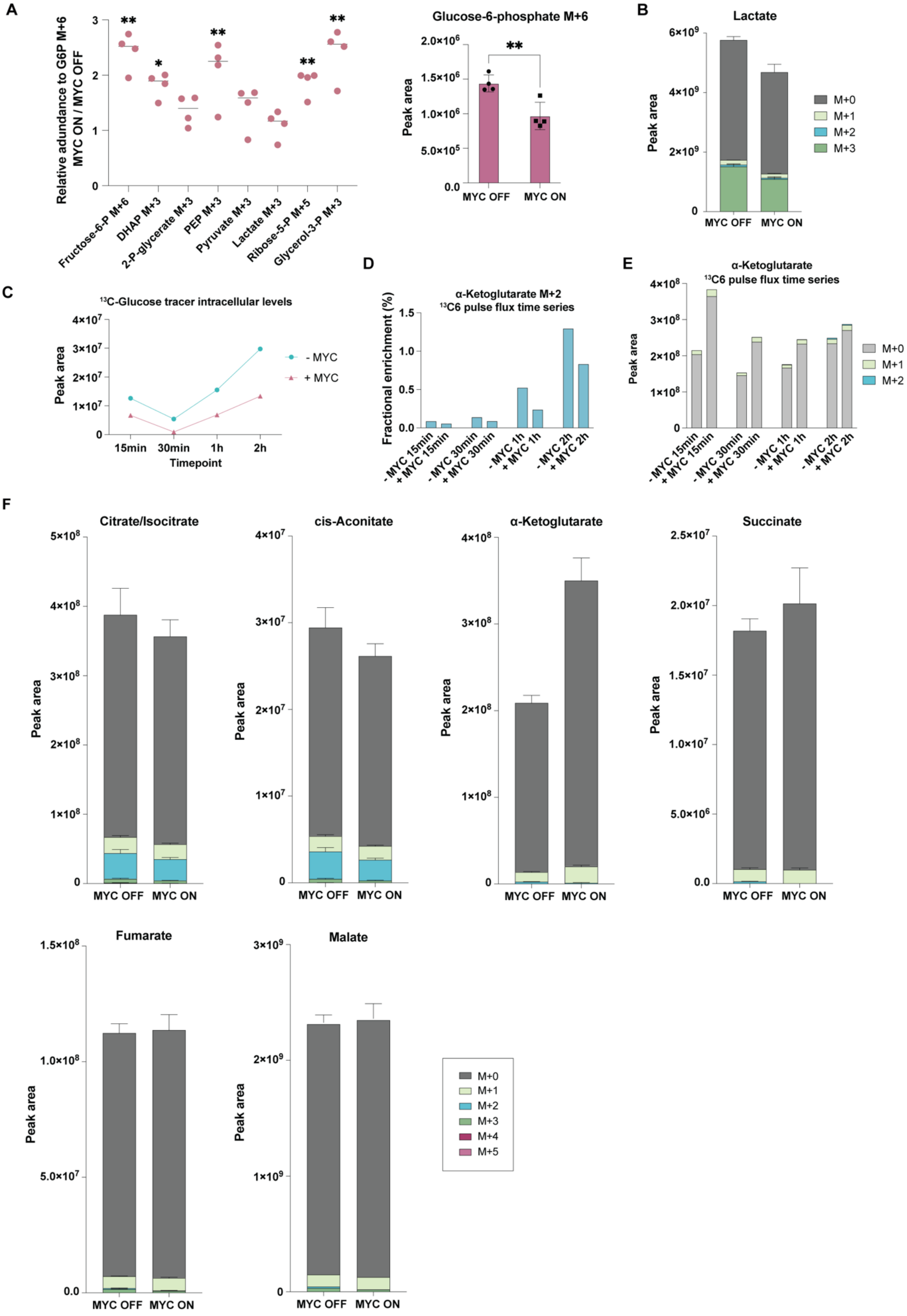

**Supplementary Figure 6. Intermediates of glycolysis, its branched pathways and TCA cycle with [U-$^{13}$C]glucose tracer. A** Relative abundance of metabolites from glycolytic pathway and its branching pathways. Each metabolite was first normalized to glucose-6-phosphate M+6 levels, followed by calculating MYC ON to MYC OFF ratio. On the right: Glucose-6-phosphate M+6 peak areas. Statistical significance: two-tailed unpaired t-test. **B** Lactate isotopologue distributions (peak areas). **C** $^{13}$C-Glucose tracer intracellular levels in time series pulse labeling experiment. N=1 for each time point. **D** Fractional enrichment of α-ketoglutarate M+2 isotopologue in the time series pulse labeling experiment. **E**. Isotopologue distribution (peak areas) of α-ketoglutarate in the time series pulse labeling experiment **F** Isotopologue distributions (peak areas) for TCA cycle intermediates citrate/isocitrate, cis-aconitate, α-ketoglutarate, succinate, fumarate and malate.

**Supplementary Figure 7.**

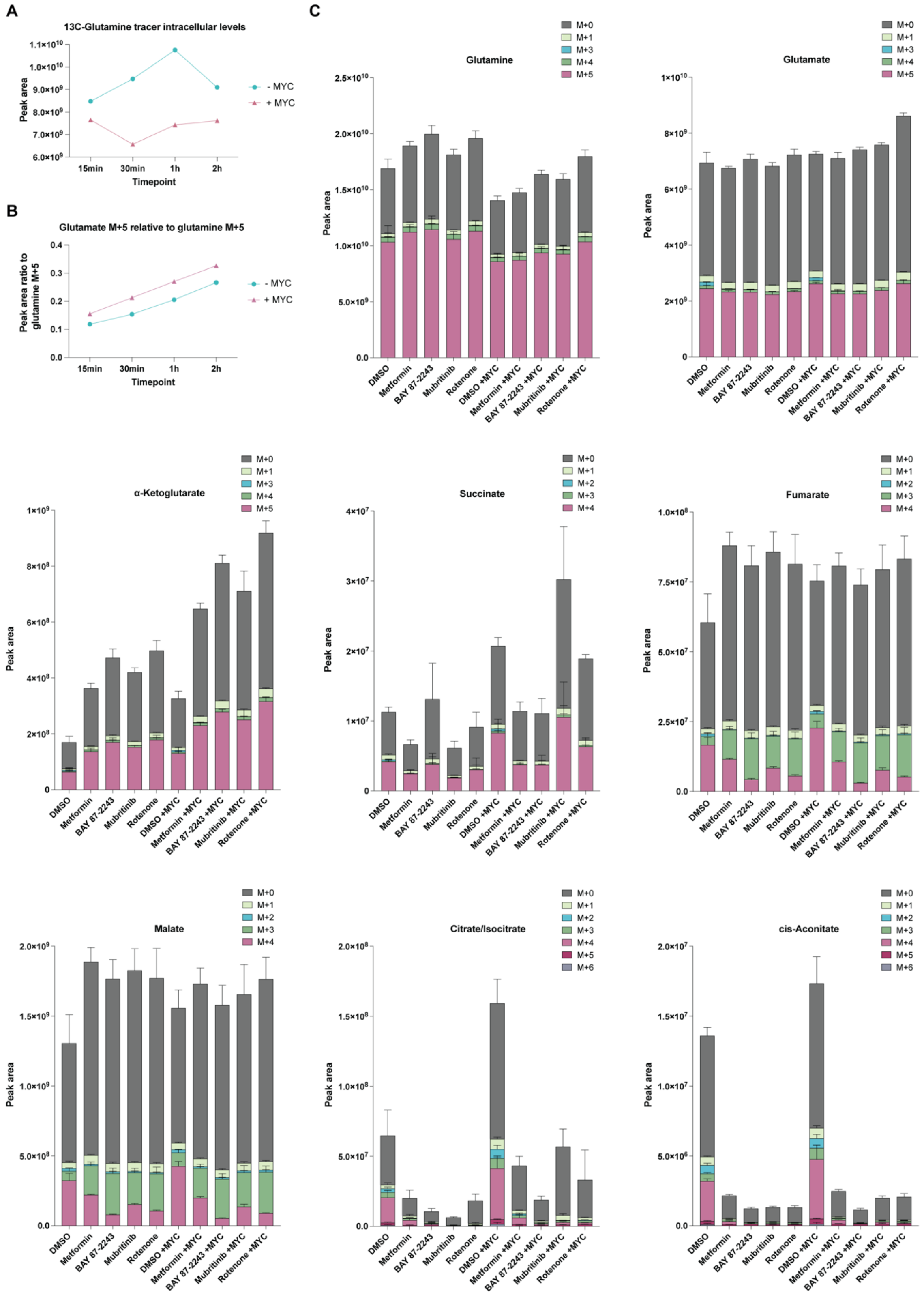


**Supplementary Figure 7. [U-$^{13}$C]glutamine tracer flux analysis of glutaminolysis and TCA cycle. A** $^{13}$C-Glutamine tracer intracellular levels in time series pulse labeling experiment. N=1 for each time point. **B** Glutamate M+5 relative levels to glutamine M+5 levels in the time series pulse labeling experiment. **C** Isotopologue distributions (peak areas) for glutamine, glutamate, α-ketoglutarate, succinate, fumarate, malate, citrate/isocitrate and cis-aconitate.

**Supplementary Figure 8.**

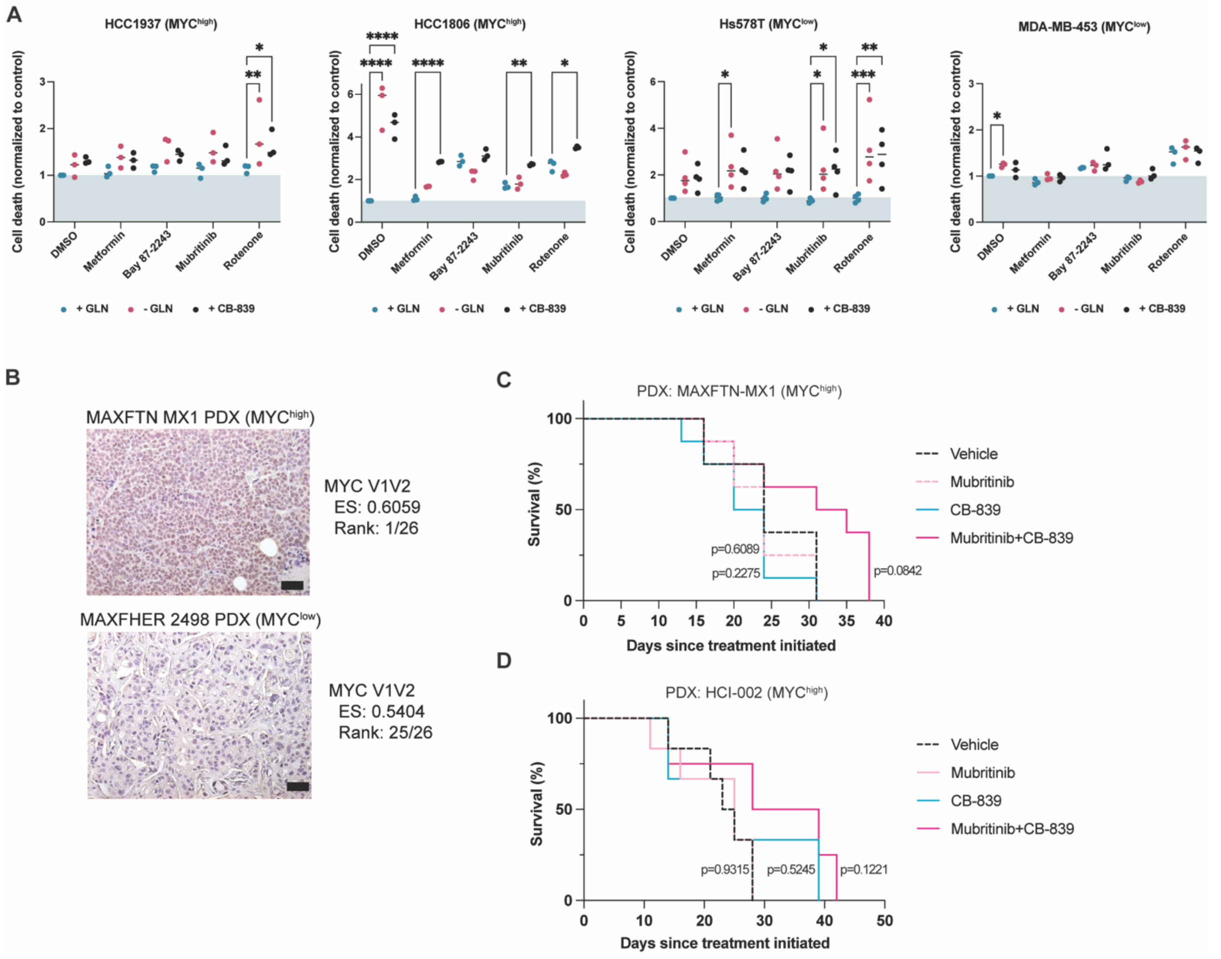


**Supplementary Figure 8. Effect of complex I inhibition and glutamine deprivation glutaminase inhibition in TNBC cell lines and MYC$^{high}$ PDX models. A** 103 CellTox Green cell death assay in MYC$^{high}$ and MYC$^{low}$ cell lines. Cells were treated with DMSO, metformin 1mM, BAY 87-2243 10 nM, mubritinib 10 nM or rotenone 30 nM. During the treatments cell were cultured in glutamine containing (+GLN), glutamine lacking (-GLN) or CB-839 containing (CB-839) medium, all in 1% FBS. Cell death was analyzed 72h after the start of treatments using IncuCyte. Statistical significance: two-way ANOVA followed by Dunnett's multiple comparisons test. **B** MYC IHC staining and MYC V1V2 signature ranking in MAXFTN MX1 (MYC$^{high}$) and MAXFHER 2496 (MYC$^{low}$) PDX models. **C** Survival of MAXFTN-MX1 PDXs. Statistical significance: Log-rank (Mantel-Cox) test. **D** Survival of HCI-002 PDXs. Statistical significance: Log-rank (Mantel-Cox) test.

**Supplementary Figure 9.**

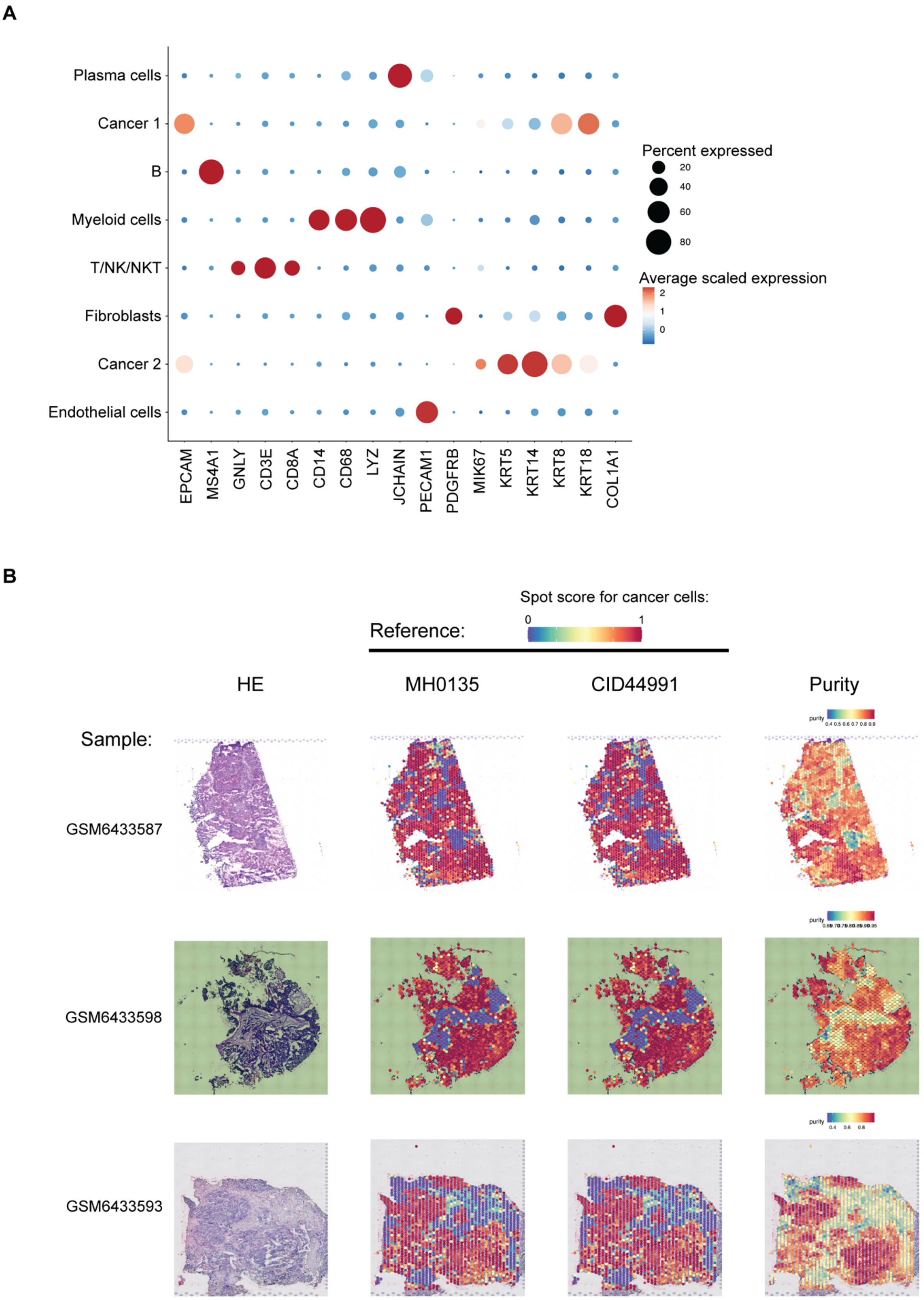


**Supplementary Figure 9. Cell type markers in scRNA-seq and tumor cell identification in spatial transcriptomics. A** Conventional markers per cell type in the scRNAseq data. **B**

Three different cell identification methods used in spatial transciptomics analysis to reliably identify tumor cell containing spots. All methods delivered very similar results, so eventually sample MH0135 was used as the reference.

Supplementary Table 1. Antibody list.

| Protein | Antibody |
|---|---|
| MYC | Ab32072, Abcam |
| H3K9me3 | Ab8898, Abcam |
| BAK | NT, 06-536, Merck Millipore |
| BAX | 6A7, sc-23959, Santa Cruz |
| pACC | Ser79, #3661, Cell Signalling Technology |
| ACC | #3676, Cell Signalling Technology |

Supplementary Data 1: RNAseq gene expression counts (provided as a separate Excel file).

Supplementary Data 2: All metabolites and their isotopogues for figure 5B.
